# Accumulation of an unprecedented 5′-deoxyadenos-4′-yl radical unmasks the kinetics of the radical-mediated C-C bond formation step in MoaA catalysis

**DOI:** 10.1101/2020.01.16.909697

**Authors:** Haoran Pang, Edward A. Lilla, Pan Zhang, Du Zhang, Thomas P. Shields, Lincoln G. Scott, Weitao Yang, Kenichi Yokoyama

## Abstract

Radical *S*-adenosyl-L-methionine (SAM) enzymes catalyze various free radical-mediated reactions. In these enzymes, the rate-determining SAM cleavage kinetically masks all the subsequent steps. Due to this kinetic masking, detailed mechanistic characterization of radical transformations catalyzed by these enzymes is very difficult. Here, we report a successful kinetic characterization of the radical C-C bond formation catalyzed by a MoaA radical SAM enzyme. MoaA catalyzes an unprecedented 3′,8-cyclization of GTP into 3′,8-cyclo-7,8-dihydro-GTP (3′,8-cH_2_GTP) during the molybdenum cofactor (Moco) biosynthesis. Through a series of EPR and biochemical characterization, we found that MoaA accumulates a 5′-deoxyadenos-4′-yl radical (5′-dA-C4′•) under the turnover conditions, and forms (4′*S*)-5′-deoxyadenosine ((4′*S*)-5′-dA), which is a C-4′ epimer of the naturally occurring (4′*R*)-5′-dA. Together with kinetic characterizations, these observations revealed the presence of a shunt pathway in which an on-pathway intermediate, GTP C-3′ radical, abstracts H-4′ atom from 5′-dA to transiently generate 5′-dA-C4′• that is subsequently reduced stereospecifically to yield (4′*S*)-5′-dA. Detailed kinetic characterization of the shunt and the main pathways provided the comprehensive view of MoaA kinetics, and determined the rate of the on-pathway 3′,8-cyclization step as 2.7 ± 0.7 s^−1^. Together with DFT calculations, this observation suggested that the 3′,8-cyclization is accelerated by 6 ∼ 9 orders of magnitude by MoaA. Potential contributions of the active-site amino acid residues, and their potential relationships with human Moco deficiency disease are discussed. This is the first determination of the magnitude of catalytic rate acceleration by a radical SAM enzyme, and provides the foundation for understanding how radical SAM enzymes achieve highly specific radical catalysis.

## Introduction

Radical *S*-adenosyl-L-methionine (SAM) enzymes^1^ are emerging group of enzymes catalyzing unique and chemically challenging reactions by free-radical mediated mechanisms. These enzymes carry out reductive cleavage of SAM using an oxygen-sensitive [4Fe-4S] cluster (Figure 1A), and transiently generate 5′-deoxyadenosyl radical (5′-dA•) to initiate radical-mediated catalysis. Despite the abundance of radical SAM enzymes and the diversity of chemical reactions they catalyze^2-3^, many fundamental aspects of the catalytic mechanisms of radical SAM enzymes remain ambiguous. It is largely unknown how these enzymes control the reactivities of radical intermediates to achieve the reaction specificity, and how much catalytic rate acceleration is provided by the enzyme active site environment. Therefore, contributions from the enzyme active site environment are frequently not considered. This significant knowledge gap is caused at least partly because all the radical chemistry is kinetically masked by the preceding slow SAM cleavage step, and thus not mechanistically tractable. Kinetic unmasking of the radical chemistry is the first critical step to understand the mechanism of radical catalysis by radical SAM enzymes. Here, using MoaA, an essential enzyme in Molybdenum cofactor (Moco) biosynthetic pathway, we report the first determination of the rate constant for the radical-mediated C-C bond formation in radical SAM enzymes. This work provides, to our knowledge, the first experimental evidence that a radical SAM enzyme accelerates a radical mediated reaction.

**Figure 1.**
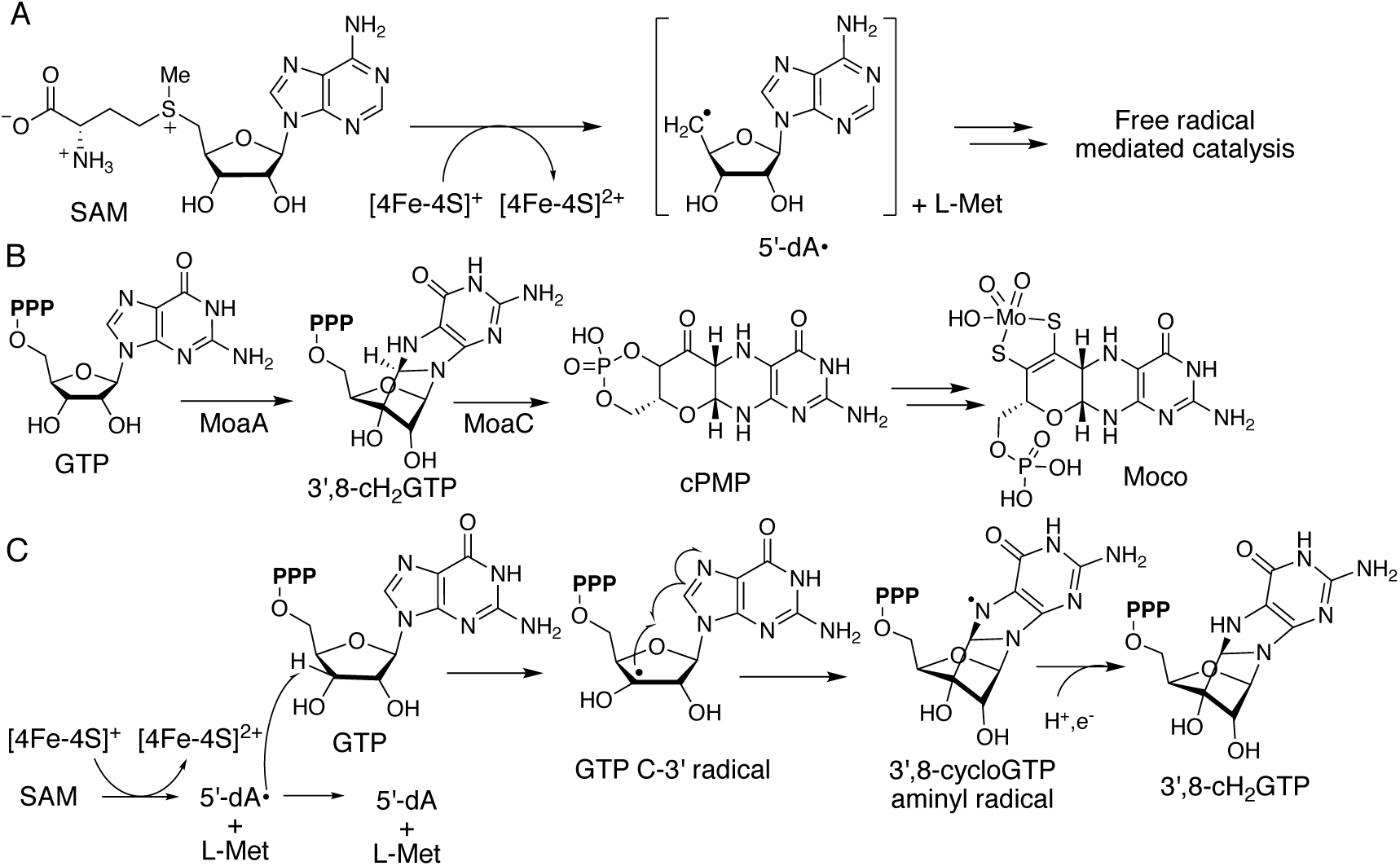
Moco biosynthetic pathway and proposed mechanism of MoaA. (A) SAM reductive cleavage catalyzed by radical SAM enzymes. (B) Moco biosynthetic pathway in bacteria. (C) Proposed mechanism of radical-mediated C3′-C8 bond formation by MoaA.

Moco is a redox enzyme cofactor found in all domains of life^4-8^. Moco-dependent enzymes play key steps in many biological pathways critical for life including purine and sulfur catabolism, anaerobic respiration, and nitrate assimilation^5^. While a salvage pathway was recently suggested in nematodes^9^, most organisms are thought to be incapable of taking up Moco as a nutrient due to the cofactor′s chemical instability, making the de novo Moco biosynthesis an essential process. In humans, Moco biosynthesis is essential for healthy development of brains. Inability to biosynthesize Moco causes fatal metabolic disorder, Moco deficiency (MoCD)^10^. In bacteria, some of the Moco-dependent enzymes are involved in the virulence of several clinically important pathogens, such as *Mycobacterium tuberculosis*^11^ and *Pseudomonas aeruginosa*^12^. In all organisms, the characteristic pterin backbone structure of Moco is biosynthesized by a set of conserved enzymes through the transformation of GTP into cyclic pyranopterin monophosphate (cPMP) (Figure 1B)^6, 13^. In bacteria, this transformation is catalyzed by two enzymes, MoaA and MoaC. MoaA catalyzes the transformation of GTP into an oxygen sensitive intermediate, 3′,8-cyclo-7,8-dihydro-GTP (3′,8-cH_2_GTP)^14^, which is then transformed into cPMP by a complex rearrangement catalyzed by MoaC (Figure 1B)^15^.

The MoaA catalysis has been proposed to proceed through a hydrogen atom abstraction from the 3′-position of GTP by 5′-dA• (Figure 1C)^14^. The resulting GTP C-3′ radical (GTP-C3′•) attacks C-8 of guanine to generate a 3′,8-cyclo-GTP aminyl radical, which is converted to 3′,8-cH_2_GTP by transfers of an electron and a proton. Among the later steps after SAM cleavage, the addition of GTP-C3′• to C-8 is likely the most chemically challenging step. Nucleotide C-3′ radicals have been reported as intermediates of several biochemical reactions, including the reactions catalyzed by ribonucleotide reductase (RNR), and an oxidative damage of DNA. In RNR, C-3′ radical facilitates dissociation of 2′-OH during the formation of deoxynucleotides^16-18^. Perturbations in the active site that prevents the completion of RNR catalysis most frequently result in base dissociation^19-21^. C-3′ radical-mediated base dissociation was also reported for oxidative damage of DNA^22^. Thus, the addition of GTP-C3′• to C-8 is an unprecedented reactivity of C3′ radical in either biological or synthetic reactions. The mechanism by which MoaA catalyzes this unprecedented C3′-C8 cyclization is unknown.

In this work, we initially aimed to kinetically and spectroscopically characterize the MoaA catalysis, which resulted in an observation of an organic radical species. Characterization of this radical by EPR using a series of site-specifically ^2^H or ^13^C-labeled SAM isotopologues revealed the observed radical as 5′-deoxyadenos-4′-yl radical (5′-dA-C4′•). When the reaction was performed with [3′-^2^H]GTP, 5′-dA-C4′• with a deuterium at the 5′ position was observed, suggesting that 5′-dA-C4′• was formed after the transfer of hydrogen from 3′-position of GTP to 5′-dA. Upon revisiting the MoaA reaction products, we observed formation of (4′*S*)-5′-dA, a 4′-epimer of (4′*R*)-5′-dA produced by the reductive cleavage of SAM. Kinetic characterization of formations of all the products and 5′-dA-C4′• and their global analysis allowed us to propose a refined model for MoaA catalysis. In this model, after abstraction of H-3′ atom from GTP by 5′-dA•, the reaction diverges to two pathways; the normal pathway that yields 3′,8-cH_2_GTP and (4′*R*)-5′-dA and a shunt pathway that yields (4′*S*)-5′-dA via 5′-dA-C4′•. This analysis also determined the rate constant for the on-pathway 3′,8-cyclization step to be 2.7 ± 0.7 s^−1^. Since DFT calculation suggested that the 3′,8-cyclization of GTP in solution without catalysis would proceed at a rate of 10^−6^ ∼ 10^−9^ s^−1^, the observed kinetics suggested that MoaA accelerates the 3′,8-cyclization by 6 ∼ 9 orders of magnitude. To our knowledge, this is the first kinetic characterization of the post-SAM cleavage reactions and the first estimation for the magnitude of rate acceleration for a radical reaction catalyzed by a radical SAM enzyme. These observations provide the foundation to understand the mechanism by which MoaA catalyzes the chemically challenging 3′,8-cyclization, and contribute to our understanding of the mechanism by which radical SAM enzymes catalyze highly specific radical reactions.

## Results

### Observation of an organic radical species in MoaA reaction

To better elucidate the mechanism of MoaA catalysis, we initially characterized wt-MoaA using X-band EPR spectroscopy. The anaerobically purified and reconstituted MoaA carried 5.8 ± 0.6 Fe per monomer, corresponding to 1.45 ± 0.15 eq. of 4Fe-4S cluster per monomer. In the absence of substrates, the EPR spectrum of MoaA at 15 K showed features that can be simulated as a sum of two axial signals (Figure 2A). The narrower signal was designated as species 1, and the wider signal as species 2. The *g* values and the ratio of the two signals are listed in Table 1. The ratio of the two axial signals and the total spin concentration were not affected by the presence of SAM, but the presence of GTP slightly increased the amount of reduced clusters (from 0.94 ± 0.09 eq. to 1.11 ± 0.06 eq. per monomer) and significantly increased the population of the species 2 (Figure 2B and 2C, Table 1). The two signals were not affected by the nature of the buffer systems (Tris, HEPES, or triethanolamine; Figure S1), and thus suggest the presence of two [4Fe-4S]^+^ cluster species whose ratio shifts in a GTP-dependent manner. The nature of the two species is currently unknown.

**Table 1:** The simulated *g* values and ratio of the two axial signals from the reduced [4Fe-4S] clusters. Total amounts of reduced [4Fe-4S] clusters are quantified by EPR based on 1 mM Cu standard. Shown are averages of three replicates and the errors represent standard deviation.

|  | substrate | species 1 |  | species 2 |  | species 1 % | species 1:2 | total [4Fe-4S]<sup>+</sup><br>(eq. of MoaA) |
| --- | --- | --- | --- | --- | --- | --- | --- | --- |
| | | $g_{\perp}$ | $g_{\parallel}$ | $g_{\perp}$ | $g_{\parallel}$ | | | |
| MoaA<br>[4Fe-4S]<sup>+</sup> | — | 1.897 | 2.031 | 1.875 | 2.063 | 57.4 ± 1.5 | 1.35 ± 0.08 | 0.94 ± 0.09 |
|  | SAM | 1.897 | 2.031 | 1.875 | 2.063 | 56.0 ± 1.5 | 1.28 ± 0.08 | 0.94 ± 0.07 |
|  | GTP | 1.890 | 2.029 | 1.881 | 2.063 | 23.1 ± 1.0 | 0.30 ± 0.02 | 1.11 ± 0.06 |
|  | GTP + SAM | 1.890 | 2.029 | 1.881 | 2.063 | 24.1 ± 1.2 | 0.32 ± 0.02 | 1.07 ± 0.01 |

**Figure 2.**
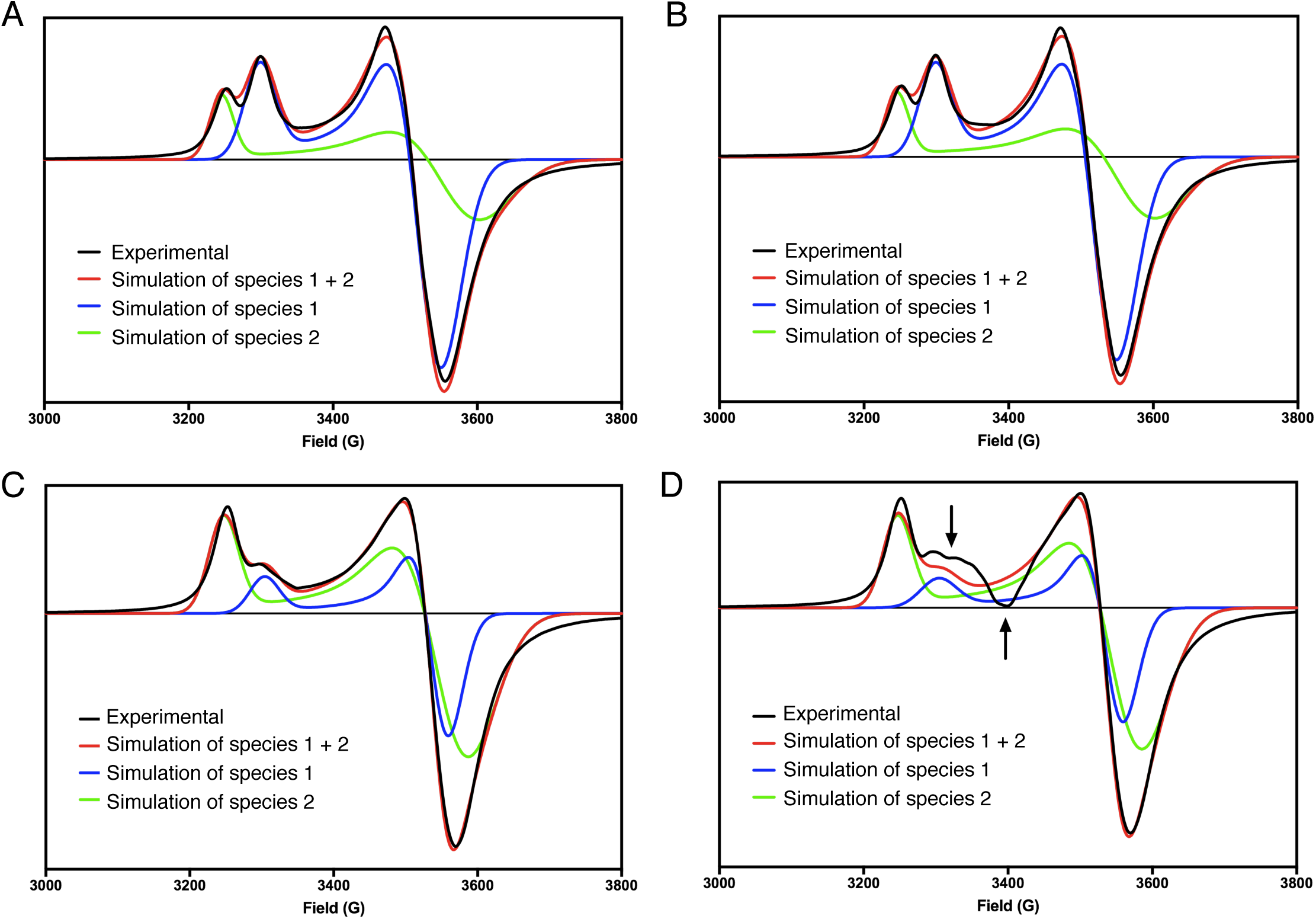
EPR spectra of MoaA at 15 K. Pre-reduced MoaA was incubated with (A) buffer only, (B) SAM, (C) GTP, (D) both SAM and GTP for 2 min and manually frozen in an isopentane slush bath.

Remarkably, in the presence of both GTP and SAM, an additional feature at g = 2.0 was observed (Figure 2D). When the EPR spectra were determined between 15 ∼ 90 K, the new feature became better resolved as the temperature increases from 15 to 45 K (Figure 3A). The intensity of this signal decreased at temperatures above 55 K. The observed temperature dependent behavior of this signal suggested that it is likely an organic radical with a fast spin relaxation property.

**Figure 3.**
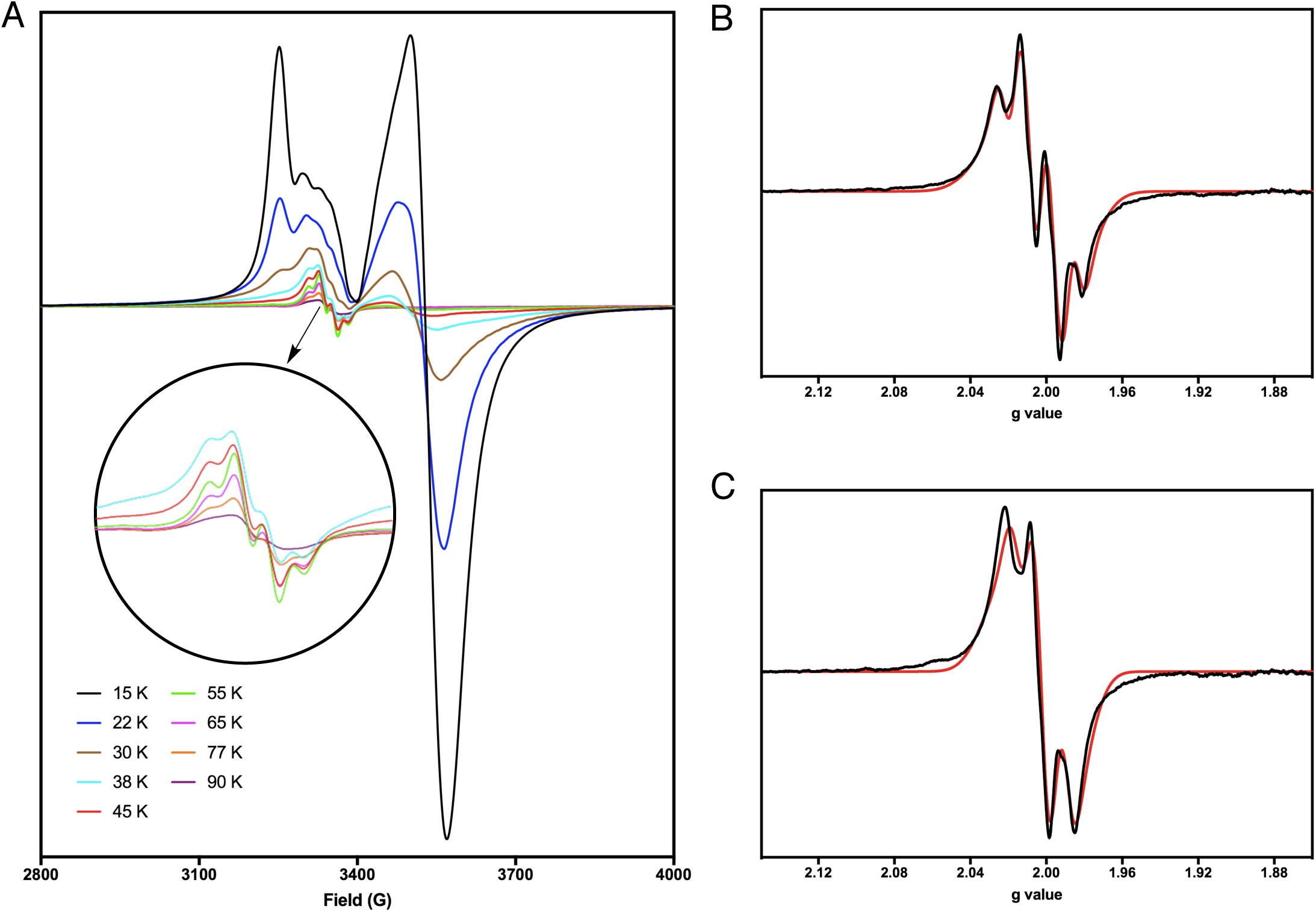
EPR characterization of the observed radical species. (A) Overlay of EPR spectra of MoaA under turnover conditions. The EPR spectra were determined at the temperatures shown in the figure. (B, C) Experimental (black trace) and simulated EPR spectra (red trace) of radical X (B) and Y (C) in MoaA reaction with GTP or [3′-^2^H]GTP as substrate, respectively. The EPR spectra were determined at 45 K and the residual [4Fe-4S]^+^ cluster signal was subtracted using a negative control without SAM.

At 45 K, the spectrum had approximately ∼ 100 G width centered at *g* = 2.0, consistent with an organic radical (Figure 3B, black trace). Intriguingly, we observed a distinct EPR spectrum when [3′-^2^H]GTP was used as substrate (Figure 3C, black trace). We hereby tentatively designate the signal observed with GTP with natural isotope abundance as X, and the signal with [3′-^2^H]GTP as Y. Radical X was simulated with hyperfine coupling to three *S* = 1/2 nuclei, suggesting the presence of three protons adjacent to the radical (Figure 3B, red trace, Table 2). Radical Y spectrum was simulated by parameters identical to X, but with one of the three *S* = 1/2 nuclei replaced with a *S* = 1 nucleus (Figure 3C, red trace, Table 2). These observations suggested that X and Y represent the same radical species with different hyperfine splitting patterns due to the replacement of one of the three ^1^H with ^2^H.

**Table 2:** The *g* values and hyperfine tensors of all the simulated spectra of 5′-dA-C4′•.

| Radical | GTP | SAM | $g_{xx}$ | $g_{yy}$ | $g_{zz}$ | $A1_{xx}$ | $A1_{yy}$ | $A1_{zz}$ | $A2_{xx}$ | $A2_{yy}$ | $A2_{zz}$ | $A3_{xx}$ | $A3_{yy}$ | $A3_{zz}$ | $A4_{xx}$ | $A4_{yy}$ | $A4_{zz}$ | lwpp (mT) |
| --- | --- | --- | --- | --- | --- | --- | --- | --- | --- | --- | --- | --- | --- | --- | --- | --- | --- | --- |
| X | GTP | SAM | 2.0059 | 2.0068 | 2.0092 | 72.1 | 66.6 | 57.6 | 43.7 | 66.1 | 113.6 | 64.1 | 21.3 | 111.3 | — | — | — | 1.07 |
|  |  | [5'-<sup>13</sup>C]SAM | 2.0059 | 2.0068 | 2.0092 | 72.1 | 66.6 | 57.6 | 43.7 | 66.1 | 113.6 | 64.1 | 21.3 | 111.3 | 19.4 | 28.0 | 31.2 | 1.08 |
|  |  | [5'-<sup>2</sup>H<sub>2</sub>]SAM | 2.0059 | 2.0068 | 2.0092 | 72.1 | 66.6 | 57.6 | 6.7 | 10.2 | 17.5 | 9.9 | 3.3 | 17.1 | — | — | — | 1.08 |
|  |  | [4'-<sup>13</sup>C]SAM | 2.0059 | 2.0068 | 2.0092 | 72.1 | 66.6 | 57.6 | 43.7 | 66.1 | 113.6 | 64.1 | 21.3 | 111.3 | 42.1 | 215.3 | 202.4 | 2.23 |
|  |  | [ribose-<sup>13</sup>C<sub>5</sub>]SAM | 2.0059 | 2.0068 | 2.0092 | 72.1 | 66.6 | 57.6 | 43.7 | 66.1 | 113.6 | 64.1 | 21.3 | 111.3 | 42.1 | 215.3 | 202.4 | 2.19 |
| Y | [3'-<sup>2</sup>H]GTP | SAM | 2.0059 | 2.0068 | 2.0092 | 11.1 | 10.2 | 8.9 | 43.7 | 66.1 | 113.6 | 64.1 | 21.3 | 111.3 | — | — | — | 1.08 |
|  |  | [5'-<sup>13</sup>C]SAM | 2.0059 | 2.0068 | 2.0092 | 11.1 | 10.2 | 8.9 | 43.7 | 66.1 | 113.6 | 64.1 | 21.3 | 111.3 | 19.4 | 28.0 | 31.2 | 1.07 |
|  |  | [5'-<sup>2</sup>H<sub>2</sub>]SAM | 2.0059 | 2.0068 | 2.0092 | 11.1 | 10.2 | 8.9 | 6.7 | 10.2 | 17.5 | 9.9 | 3.3 | 17.1 | — | — | — | 1.08 |
|  |  | [4'-<sup>13</sup>C]SAM | 2.0059 | 2.0068 | 2.0092 | 11.1 | 10.2 | 8.9 | 43.7 | 66.1 | 113.6 | 64.1 | 21.3 | 111.3 | 42.1 | 215.3 | 202.4 | 2.22 |
|  |  | [ribose-<sup>13</sup>C<sub>5</sub>]SAM | 2.0059 | 2.0068 | 2.0092 | 11.1 | 10.2 | 8.9 | 43.7 | 66.1 | 113.6 | 64.1 | 21.3 | 111.3 | 42.1 | 215.3 | 202.4 | 2.23 |

### The observed organic radical exhibits a unique relaxation property

In general, organic radicals are best characterized at ∼77 K. However, the radicals X and Y were broadened and less intense at 77 K than at 45 K, indicating its fast relaxation property. Thus, radical X was investigated by microwave power saturation experiments. When the radical signal intensities were quantified and plotted against the root square of microwave power, no apparent signal saturation was observed up to 16 mW and only a partial saturation was observed even at the highest possible microwave power, 200 mW. The resulting data were analyzed by a non-linear curve fit to the following equation^23^ (Figure 4):

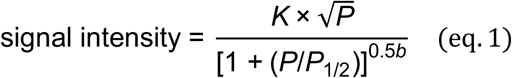

**Figure 4.**
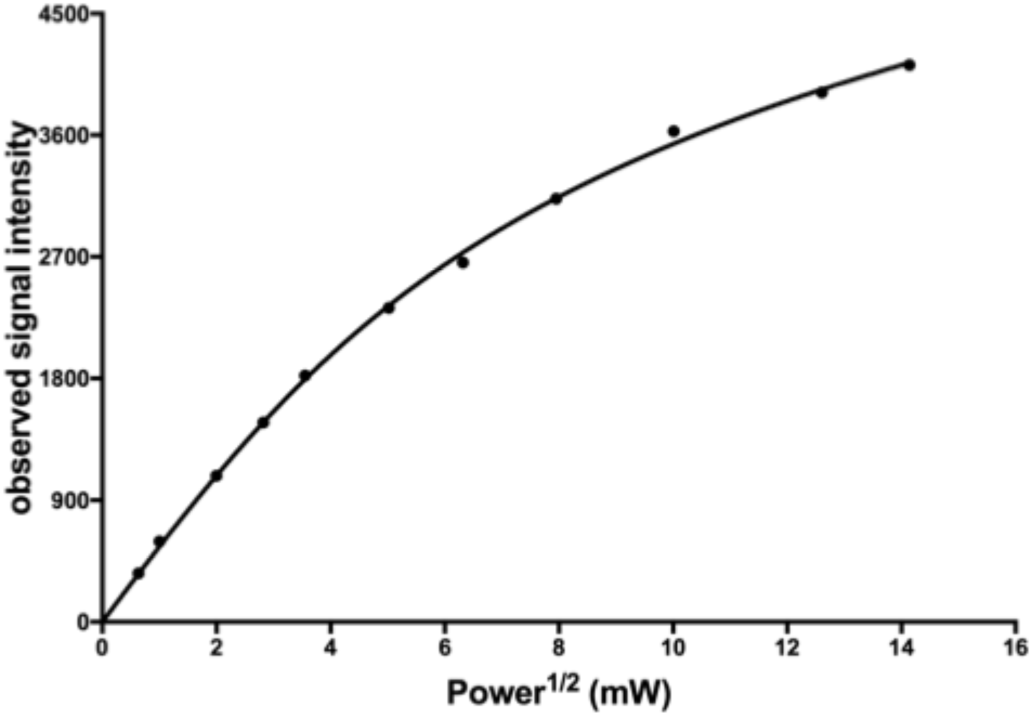
Relaxation property of radical X measured at 45 K. The solid line represents a non-linear curve fit to eq. 1 with *K* = 561.5 ± 15.4, *P*_1/2_ = 34.5 ± 10.8 mW, and *b* = 0.68 ± 0.08.

*K* is a scaling factor, *P* represents microwave power in milliwatts and *P*_1/2_ is the microwave power at half saturation, and *b* is the inhomogeneity parameter^24^. The analysis revealed *P*_1/2_ = 34.5 ± 10.8 mW with *b* = 0.68 ± 0.08. The observed *P*_1/2_ was significantly larger than those reported for organic radical without any other paramagnetic centers. In the radical SAM enzyme PolH, the substrate radical trapped in a C209A mutant enzyme active site exhibited *P*_1/2_ = 23.3 ± 17.7 μW at 77 K (Figure S1)^25^. Similarly, characterization of a substrate radical observed in Cfr requires a low microwave power (20 μW) to avoid saturation^26^. In BtrN, another member belongs to SPASM/Twitch family, a substrate radical was characterized with a microwave power of 400 μW^27^ and both [4Fe-4S] clusters in the active site were proposed to be 2+ state when the radical was formed^28^. On the other hand, in lysine 2,3-aminomutase (LAM), one of the substrate radicals observed was reported to have an onset of saturation at > 5 mW at 77 K^29^. The fast relaxation feature of radicals in LAM was attributed to the presence of another paramagnetic center although its identity is not yet known^30^. Based on all these reported observations, the relaxation property of the radical observed in MoaA was faster than any other radicals observed in other radical SAM enzymes, and suggests the presence of another paramagnetic center in the active site, most likely the reduced auxiliary [4Fe-4S] cluster.

### Characterization of the radical using isotopically labeled SAM

The hyperfine splitting pattern of signal X suggested the presence of three protons adjacent to the radical, which eliminates the possibility of GTP C-3′ or N-7 radical. Thus, we investigated the possibility of a radical on 5′-dA using isotopically labeled SAM. SAM isotopologues were synthesized enzymatically from L-methionine and site-specifically isotope labeled ATP using recombinant SAM synthetase. The isotope labeled ATP was prepared from either glucose or ribose with corresponding isotope labeling using de novo purine biosynthetic enzymes^31-32^. To investigate if the radical is indeed on 5′-dA, we first characterized radical X using [ribose-^13^C_5_]SAM. The resulting EPR spectrum showed significantly broader features with ∼ 200 G width (Figure 5A, trace 2) with a large hyperfine coupling constant (∼ 200 MHz, Table 2) consistent with a hyperfine coupling with ^13^C. These results suggest that radical X is on ribose carbon(s) of 5′-dA.

**Figure 5.**
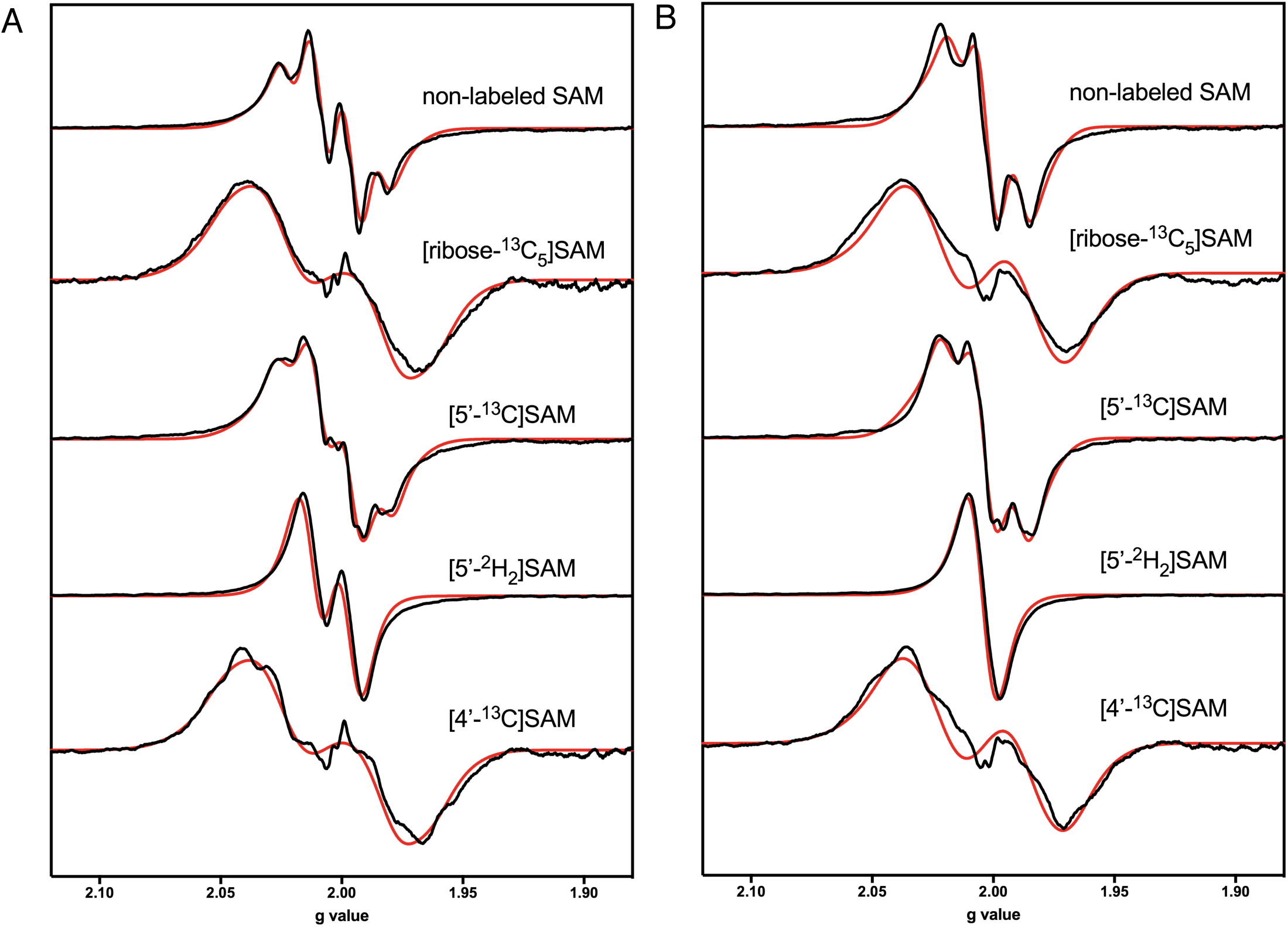
EPR characterization of radical X and Y by using isotopically labeled SAM. Shown are experimental (black traces) and simulated EPR spectra (red traces) of radicals X (A) and Y (B) generated in the presence of isotopically labeled SAM.

To determine the exact location of radical X, we prepared SAM isotopologues with either ^13^C or ^2^H on 4′- or 5′-position. When [5′-^13^C]SAM was used in the reaction, only a small hyperfine coupling constant (∼ 30 MHz) derived from the 5′-^13^C labeling was observed (Figure 5A trace 3, Table 2), indicating that the radical was not on C-5′. When [5′-^2^H_2_]SAM was used, two of the three major hyperfine splitting features in the EPR spectrum of radical X collapsed, resulting in a signal with only one apparent hyperfine splitting (Figure 5A, trace 4). The simulation revealed that the radical was hyperfine coupled to only one *S* = 1/2 nucleus (Table 2). These observations suggest that radical X is hyperfine coupled to H-5′ of SAM. When [4′-^13^C]SAM was used, the EPR spectrum of the radical exhibited a large hyperfine splitting (∼ 200 MHz) comparable to those observed in the reaction with [ribose-^13^C_5_]SAM (Figure 5A trace 5, Table 2). These observations are most consistent with radical X being a 5′-deoxyadenos-4′-yl radical (5′-dA-C4′•).

The radical Y was also characterized using the isotopically labeled SAM, which paralleled the observations with X (Figure 5B, Table 2). While [ribose-^13^C_5_]SAM caused a large hyperfine splitting (∼200 MHz), [5′-^13^C]SAM resulted in a small hyperfine coupling splitting (∼ 30 MHz), indicating that radical Y is not on C-5′. When [5′-^2^H_2_]SAM was used, the original two major hyperfine splitting features both collapsed, resulting in a singlet signal without any defined hyperfine splitting. When [4′-^13^C]SAM was used, a large hyperfine splitting (∼ 200 MHz) comparable to that with [ribose-^13^C_5_]SAM was observed. All these observations suggest that radical Y is also on C-4′. The difference in the splitting patterns of radicals X and Y likely derives from the isotope at the 3′-position of GTP. Since the ^1^H or ^2^H-atom at the 3′-position of GTP is transferred to the 5′-position of 5′-dA during the catalysis, the presence of hyperfine splitting with only two protons in radical Y can be explained by the presence of one ^2^H at the 5′-position of 5′-dA. On the basis of these considerations, we propose radical Y as [5′-^2^H]5′-dA-C4′•.

### Detection of (4′S)-5′-deoxyadenosine ((4′S)-5′-dA) formation

Based on the observation of 5′-dA-C4′•, we sought for shunt products that may be derived from 5′-dA-C4′•. When MoaA reaction was quenched by acid and analyzed by HPLC, we observed a small peak with retention time ∼2 min earlier than 5′-dA formed from SAM cleavage (Figure 6A). The UV absorption spectrum of this peak was consistent with an adenine-containing molecule. The amount of this species increased with longer reaction time. The same product was observed in both MoaA reaction or MoaA/MoaC coupled reaction (Figure 6A). To determine its identity, the minor species and 5′-dA were purified by HPLC, and individually analyzed by LCMS. The two products were eluted with different retention time on LCMS, but showed an identical molecular weight (Figure 6B). Based on these observations and in light of the existence of 5′-dA-C4′•, the minor species was assigned as (4′*S*)-5′-dA, a 4′-epimer of naturally occurring (4′R)-5′-dA. MoaA reactions with [3′-D]GTP also produced both (4′*R*)-5′-dA and (4′*S*)-5′-dA with +1 Da higher molecular weight (Figure S2A), demonstrating the (4′*S*)-5′-dA was formed after H-3′ atom abstraction from GTP by 5′-dA•. These observations suggest that (4′*S*)-5′-dA was formed by a reductive quenching of 5′-dA-C4′• by abstracting a nearby hydrogen atom or by transfers of a solvent exchangeable proton and an electron. When the MoaA reaction was conducted in D_2_O, the molecular weights of (4′*R*)-5′-dA and (4′*S*)-5′-dA were identical to those in H_2_O (Figure S2B), suggesting (4′*S*)-5′-dA is formed by 5′-dA-C4′• abstracting a solvent non-exchangeable H-atom. These results indicate that 5′-dA-C4′• is likely a radical intermediate on a shunt pathway that is subsequently quenched to yield (4′*S*)-5′-dA.

**Figure 6.**
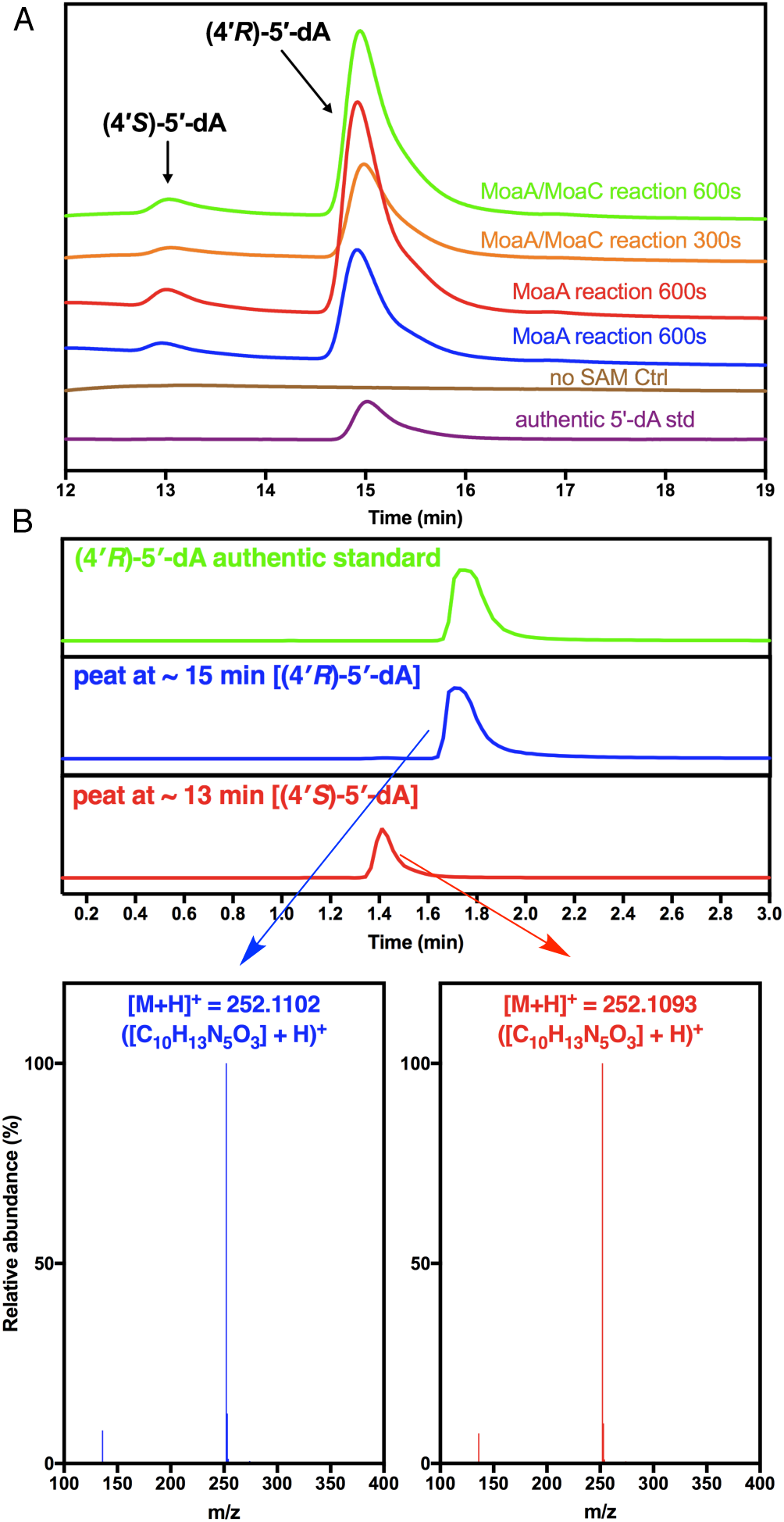
Formation of (4′*S*)-5′-dA in MoaA reaction. (A) HPLC detection of (4′*S*)- and (4′*R*)-5′-dA in MoaA assays. Shown are HPLC traces monitored at 260 nm for (from the bottom to the top) authentic (4′*R*)-5′-dA standard, a control MoaA assay without SAM, MoaA assay quenched at 300 or 600 s, MoaA/MoaC coupled assay quenched at 300 or 600 s. (B) LCMS characterization of (4′*S*)- and (4′*R*)-5′-dA isolated from the MoaA reaction with GTP and SAM. Shown are extracted ion chromatograms monitored at m/z = 252.1091, and extracted mass spectrum at each retention time.

### Kinetic characterization of MoaA catalysis

To establish a kinetic model for MoaA catalysis, the kinetics of formations of 5′-dA-C4′•, 3′,8-cH_2_GTP, (4′*R*)-5′-dA and (4′*S*)-5′-dA at pre-steady state were investigated. In these analyses, MoaA assays were performed with 300 μM MoaA, 1 mM SDT, 1 mM GTP and 1 mM SAM for 15 ∼ 300 s. To quantify 5′-dA-C4′•, the reactions manually freeze-quenched by isopentane slush bath, characterized by EPR, and quantified based on flavodoxin semiquinone standard^33-34^ (Figure S3). The time course of 5′-dA-C4′• formation (Figure 7A,B) revealed that 5′-dA-C4′• was formed in the first 2 min with the maximum amount of 1.6 ± 0.3% of MoaA (4.7 ± 1.0 μM with 300 μM MoaA). To quantify 3′,8-cH_2_GTP, (4′*R*)-5′-dA and (4′*S*)-5′-dA, the reactions were quenched by acid and analyzed by HPLC. For accurate quantitation of chemically unstable 3′,8-cH_2_GTP, MoaC was included in the assay to convert 3′,8-cH_2_GTP to cPMP, which is subsequently converted to a fluorescent derivative, compound Z (CmdZ)^14, 35^. The presence of MoaC did not affect 5′-dA-C4′• formation (Figure S4). The lag phase caused by the coupled assay was estimated to be < 0.02 s based on Storer and Cornish-Bowden′s method^36-37^. A residual amount of 3′,8-cH_2_GTP, likely due to its binding to MoaA, was quantified by derivatization to dimethylpterin (DMPT)^14, 38^. Thus, the amount of 3′,8-cH_2_GTP was a sum of CmdZ and DMPT (Figure 7A,B). (4′*R*)-5′-dA and (4′*S*)-5′-dA were quantified by HPLC with authentic (4′*R*)-5′-dA as a standard (Figure 7A and B). As shown in Fig. 7A, there was a small but statistically significant delay in the formation of 3′,8-cH_2_GTP relative to that of (4′*R*)-5′-dA in the first ∼2 min. The formation of (4′S)-5′-dA was significantly slower than that of 5′-dA-C4′• with the lag phase in the first ∼ 90 sec, but progressed over time and the amount of (4′S)-5′-dA exceeded that of 5′-dA-C4′• after 3 min (Figure 7B).

**Table 3:**
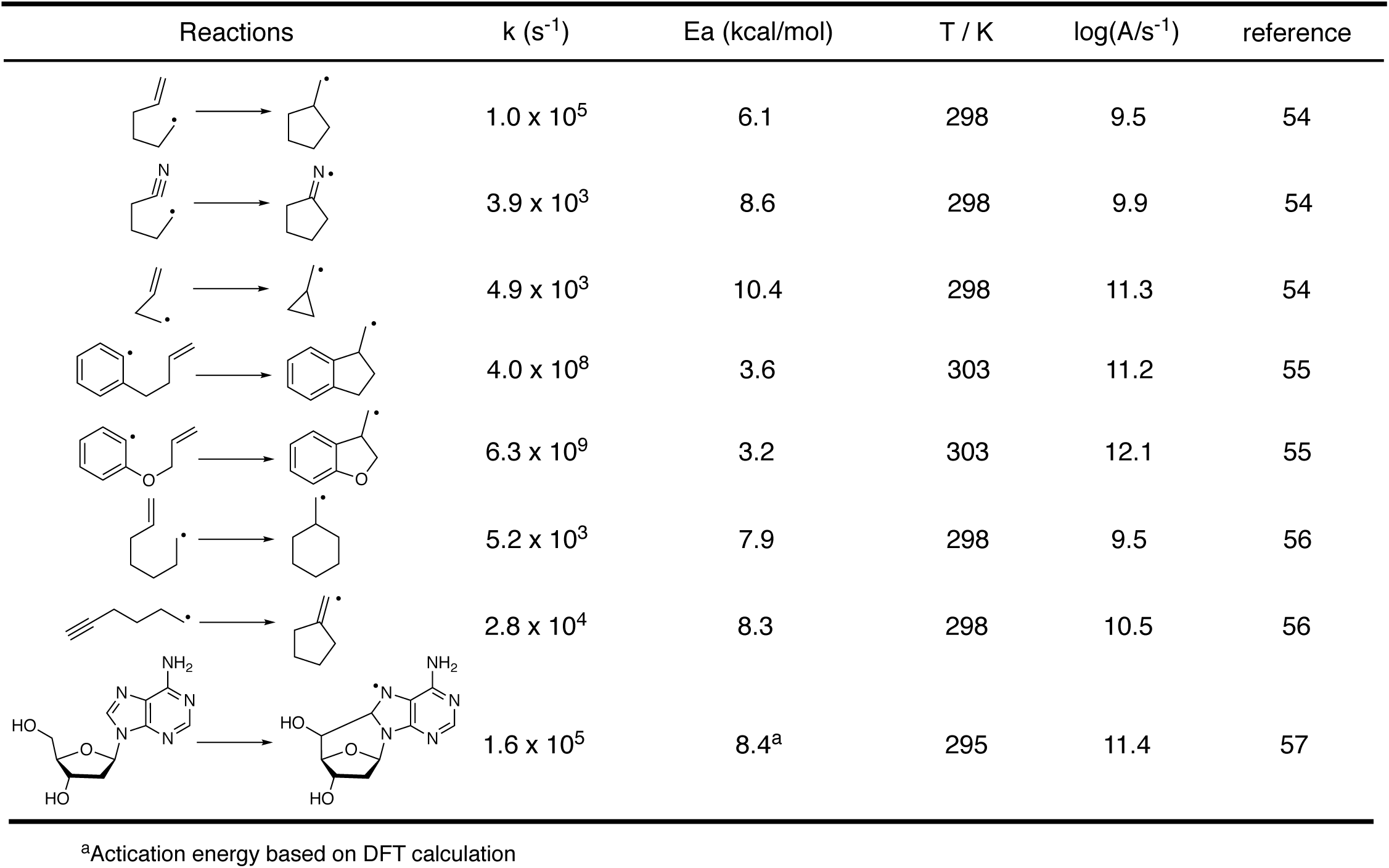
Reported rate constants and activation energies for intramolecular cyclization reactions in solution.

| Reactions | k (s<sup>-1</sup>) | Ea (kcal/mol) | T / K | log(A/s<sup>-1</sup>) | reference |
| --- | --- | --- | --- | --- | --- |
|  | 1.0 x 10<sup>5</sup> | 6.1 | 298 | 9.5 | 54 |
|  | 3.9 x 10<sup>3</sup> | 8.6 | 298 | 9.9 | 54 |
|  | 4.9 x 10<sup>3</sup> | 10.4 | 298 | 11.3 | 54 |
|  | 4.0 x 10<sup>8</sup> | 3.6 | 303 | 11.2 | 55 |
|  | 6.3 x 10<sup>9</sup> | 3.2 | 303 | 12.1 | 55 |
|  | 5.2 x 10<sup>3</sup> | 7.9 | 298 | 9.5 | 56 |
|  | 2.8 x 10<sup>4</sup> | 8.3 | 298 | 10.5 | 56 |
|  | 1.6 x 10<sup>5</sup> | 8.4<sup>a</sup> | 295 | 11.4 | 57 |

<sup>a</sup>Activation energy based on DFT calculation

**Figure 7:**
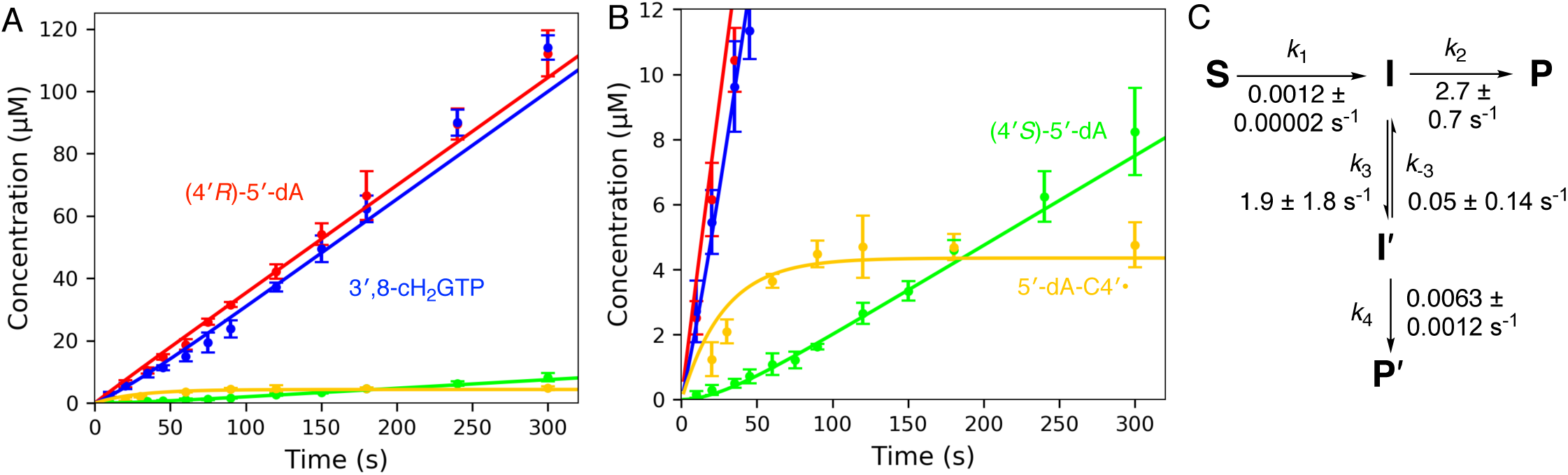
Kinetic model of MoaA catalysis. (A and B) Time course of formations of 5′-dA-C4′•, 3′,8-cH_2_GTP, (4′*R*)-5′-dA and (4′*S*)-5′-dA. B is a magnified view of A focused on 5′-dA-C4′• and (4′*S*)-5′-dA. The solid lines are the results of global fitting (p-value of 0.063 and χ^2^/DoF of 1.32) using KinTek Explorer^63-64^ and the parameters and the model shown in C. (C) Kinetic model and rate constants used to analyze the time course data in A and B. **S**, the Michaelis complex (MoaA•GTP•SAM); **I**, GTP-C3′•; **P**, 3′,8-cH_2_GTP and (4′*R*)-5′-dA; **I**′, 5′-dA-C4′•, and **P**′, (4′*S*)-5′-dA.

The kinetic data of all four species were globally fit to kinetic models, among which the shunt pathway model (Figure 7C) gave the best fit. In this model, the MoaA catalysis is consisted of two paths. The main path produces 3′,8-cH_2_GTP and (4′*R*)-5′-dA via GTP-C3′ radical, and represented by **S** → **I** → **P**, where **S** represents MoaA in complex with GTP and SAM (Michaelis complex), **I** represents GTP-C3′•, and **P** represents 3′,8-cH_2_GTP. The shunt pathway that yields (4′*S*)-5′-dA via 5′-dA-C4′• is represented by **I** → **I**′ → **P**′, where **I**′ represents 5′-dA-C4′•, and **P**′ represents the shunt product (4′*S*)-5′-dA. The concentration of the Michaelis complex (**S**) was set as an invariable at 300 μM based on the low *K*_d_ values (3 ∼ 5 μM) and excess amounts (1 mM) of SAM and GTP used in all the assays relative to MoaA (300 μM). The amount of **I** was set below the EPR detection limit (0.2 μM) because no detectable amount of GTP-C3′• was observed in our EPR characterizations.

Reversibility of each step was determined by either the kinetic fitting or available biochemical evidences. The reversibility of SAM cleavage and subsequent H-atom abstraction can be probed by the number of deuterium atoms incorporated into 5′-dA from the substrate. While multiple deuterium atoms were found in 5′-dA formed by radical SAM enzymes that catalyze reversible SAM cleavage and H-atom abstraction^39-41^, MoaA produced only singly deuterated 5′-dA from [3′-D]GTP^14^. Therefore, **S** → **I** was set irreversible. The product formation steps (**I** → **P, I**′ → **P**′) were also set irreversible because it is unlikely that the radicals are reversibly generated from these products. The reversibility of 5′-dA-C4′• formation from GTP-C3′• is unknown and both possibilities were investigated (Figure 7C, Figure S5A). Irreversible 5′-dA-C4′• formation model (Figure S6F) resulted in poorer fit compared to the reversible model (Figure S6F, Fig. 7C; p values 1.2 × 10^−10^ vs 0.063), suggesting that the data is more consistent with the reversible mechanism. However, the error values for *k*_3_ and *k*_-3_ were large, likely due to that GTP-C3′• is unquantifiable, leaving some ambiguity about the kinetic nature of the transformation between **I** and **I′**.

The acid quenching used for 5’-dA detection should have quenched the accumulated 5′-dA-C4′• to (4′*R*)-5′-dA, (4′*S*)-5′-dA or other uncharacterized molecules. Therefore, we performed kinetic fitting for cases where the acid-quenching of 5′-dA-C4′• yielded only (4′*R*)-5′-dA (Figure 7), only (4′*S*)-5′-dA (Figure S5B), both (4′*R*)-5′-dA and (4′*S*)-5′-dA with a 1:1 ratio (Figure S5C) or other uncharacterized molecules (Figure S5D). Among these possibilities, the model in which the acid quenching converted 5′-dA-C4′• to (4′*R*)-5′-dA provided the best fit.

The amount of Michaelis complex (**S**) may be lower than 300 μM due to incomplete loading of [4Fe-4S] clusters. An estimation of ∼ 1.5 eq. of [4Fe-4S] clusters per monomer were reconstituted successfully based on Ferrozine assay, and ∼ 1 eq. of [4Fe-4S] clusters per monomer were reduced by SDT based on EPR characterization. Hence 150 μM (Figure S5E) or 225 μM (Figure S5F) of **S** were also tested for global fitting. Kinetic fitting with different amount of **S** only significantly change *k*_1_ which represents rate determining step of the entire reaction. While the concentration of Michaelis complex affected *k*_*1*_, the *k*_2_ and *k*_3_ values were only minimally affected (*k*_2_ = 2.5 - 2.7 s^−1^, *k*_3_ = 1.9 – 2.7 s^−1^, and *k*_3_ = 0.050 – 0.084 s^−1^).

Another potential model where reductive quenching of 5′-dA-C4′• in the shunt pathway yields not only (4′*S*)-5′-dA, but also (4′*R*)-5′-dA (Figure S5G) was considered. Kinetic fitting with inclusion of (4′*R*)-5′-dA (**A**) into the shunt pathway resulted in an extremely low rate constant for **I**′ → **A**, indicating reductive quenching of 5′-dA-C4′• to (4′*R*)-5′-dA is unlikely. This observation is consistent with the absence of incorporation of a deuterium to (4′*S*)-5′-dA from solvent (Figure S2B), which suggested a specific mechanism of 5′-dA-C4′• quenching to form (4′*S*)-5′-dA.

The rate constants in the best global fit are shown in Fig. 7C. Most significantly, the analysis allowed determination of the rate constant for the transformation of GTP-C3′• (**I**) into 3′,8-cH_2_GTP (**P**, 2.7 ± 0.7 s^−1^). Since the aminyl radical intermediate did not accumulate at the detectable amount, the GTP-C3′• addition to C8 is likely the rate determining step of this transformation, and thus represented by *k*_2_. The *k*_1_ is the rate determining step of the entire reaction and consistent with the reported *k*_cat_ value determined under steady-state conditions. The observed *k*_2_ is 3 orders of magnitude faster than *k*_1_, which is consistent with GTP-C3′• (**I**) not experimentally detectable. Therefore, the presence of the kinetically competitive shunt pathway (*k*_3_ = 1.9 ± 1.8 s^−1^) allowed the first experimental determination of the rate constant for the kinetically masked 3′,8-cyclization step. To our knowledge, this is the first determination of the rate constant of post-H-atom abstraction steps catalyzed by radical SAM enzymes.

### Evaluation of the catalytic rate acceleration

To evaluate the rate constant from kinetic fitting, we compared experimentally determined rate constants and activation energies for intramolecular radical cyclization where alkyl radicals attack sp^2^ or sp^1^ carbon centers^42-45^ (Table 3). The examples include the 5′,8-cyclization of 2′-deoxyadenosine, which is known as a cyclopurine lesion of DNA oxidative damage and has been well-characterized using model systems. The reported rate constant for this reaction was experimentally determined as 1.6 × 10^5^ s^−1 45^. While no experimental activation energy (*E*_a_) for this reaction has been reported, a theoretical value based on DFT calculation has been reported as 8.4 kcal/mol^45^. In these examples, the natural log of the pre-exponential factors (A) of the Arrhenius equation falls in a relatively narrow range of 9.5 ∼ 12.1.

Since 3′,8-cyclization does not proceed in solution without enzymatic catalysis, no experimental rate constants or *E*_a_ is available. Instead, we performed a DFT calculation of the 3′,8-cyclization in solution at 298 K. Calculation of a transformation of GTP-C3′• to 3′,8-cyclo-GTP aminyl radical in solution without any catalysis revealed a theoretical *E*_a_ of 24.6 kcal/mol (Fig. 8A). Assuming that the pre-exponential factor (A) of the Arrhenius equation for the 3′,8-cyclization is similar to those of the related radical cyclization reactions in Table 3, the *E*_a_ correspond to rate constants of 1.1 × 10^−6^ ∼ 2.9 × 10^−9^ s^−1^. These evaluations are consistent with the absence of reported 3′,8-cyclization in non-enzymatic reactions and the chemically challenging nature of this transformation. For the MoaA-catalyzed 3′,8-cyclization, based on the rate constant *k*_2_ = 2.7 ± 0.7 s^−1^ and the A value in Table 3, the *E*_a_ was estimated as 12.2 ∼ 16.1 kcal/mol. Therefore, these analyses suggest that MoaA likely stabilizes the transition state by 8.5 ∼ 12.4 kcal/mol, and accelerate the reaction rate by 6 ∼ 9 orders of magnitude.

**Figure 8.**
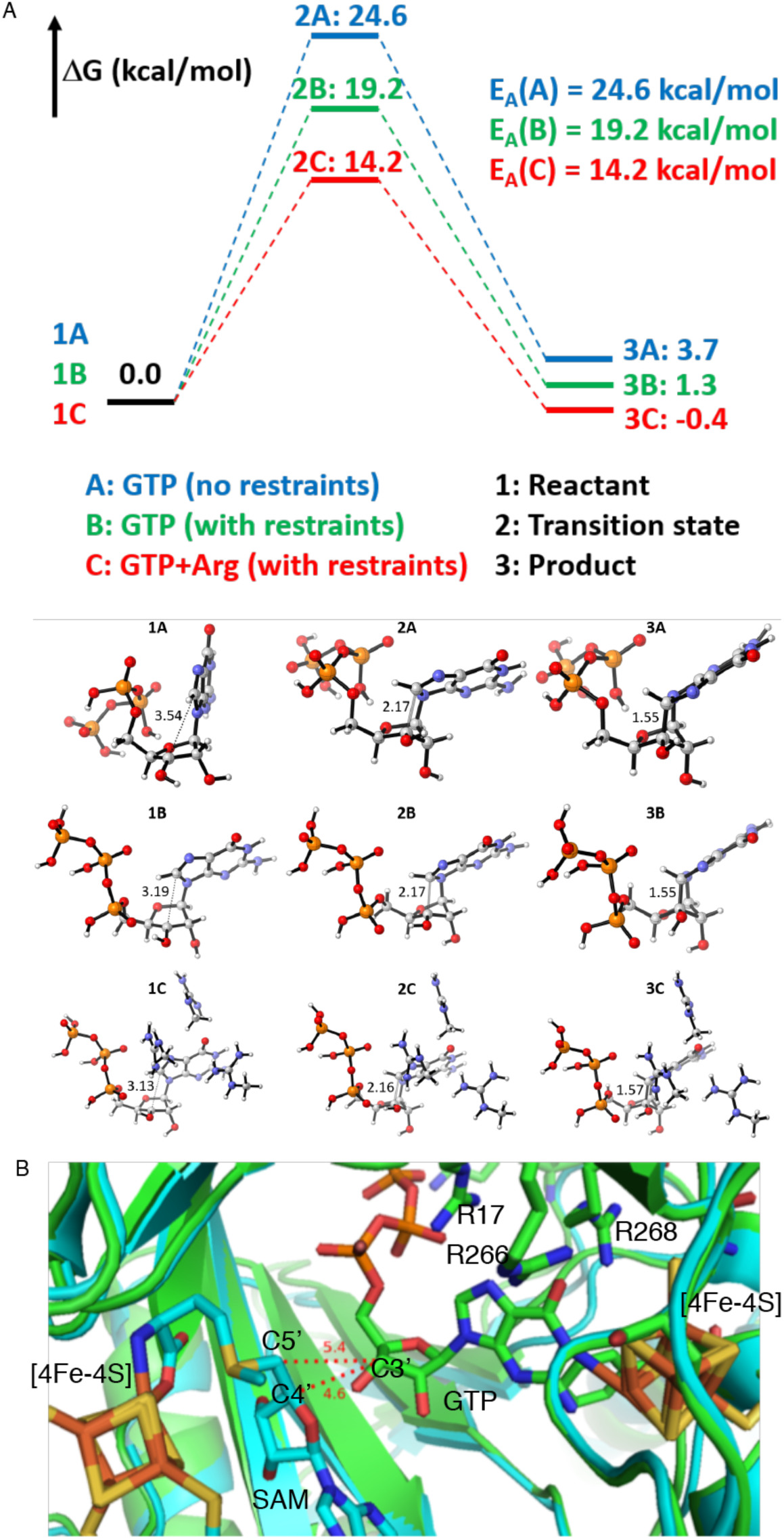
DFT calculation of 3′,8-cyclization and model of MoaA active site. (A) Reaction coordinates of C3′-C8 bond formation from GTP-C3′• to yield 3′,8-cyclo-GTP aminyl radical under three conditions: (1) in solution without any catalysis, (2) conformation constrained based on the crystal and solution structure of GTP bound to MoaA, (3) the conformationally constrained model (2) with three catalytically essential Arg residues. (B) Model of the MoaA active site is created by fusing the reported structure of MoaA in complex with SAM^65^ (PDB ID: 1TV8, cyan) and with GTP^38^ (PDB ID: 2FB3, green). Atom color: oxygens are shown in red, nitrogens in blue, phosphorus in orange, irons in brown, sulfurs in yellow.

We further investigated the effect of the active site environment on the *E*_a_ of 3′,8-cyclization. Conformational restraints were first applied to the structure of GTP-C3′•. The triphosphate moiety and O6, N1, and N2 atoms on guanine base were fixed based on the reported X-ray crystal structure^38^ and ENDOR analysis^46^ of MoaA in complex with GTP, where extensive interactions were observed between these positions of GTP and MoaA active site amino acid residues or the auxiliary cluster. The ribose moiety was not H-bonded to any of the amino acid residues, and thus allowed to move. When this partially fixed GTP-C3′• was used, the theoretical *E*_a_ of 3′,8-cyclization was calculated as 19.2 kcal/mol (Fig. 8A), which is ∼ 5.4 kcal/mol lower than the calculation without any conformational constrains. This lowering in *E*_a_ may be explained by the energy minimized conformation of GTP-C3′• (1C in Figure 8), in which C3′ is already in an orientation favorable to attack C8 with a relatively short distance between C3′ and C8 (3.19 Å, Figure 8A). Longer C3’-C8 distances were found in GTP-C3′• without any constraints (3.54 Å, 1A in Figure 8A) or in the X-ray crystal structure of MoaA•GTP complex (3.8 Å). These observations suggest that the MoaA active site likely pre-organizes the GTP-C3′• conformation to accelerate the 3′,8-cyclization.

The theoretical *E*_a_ determined for the conformational constraint model, however, was still higher than the estimated *E*_a_ of 12.2 ∼ 16.1 kcal/mol. Therefore, we investigated the effects of the three catalytically essential Arg residues (R17, R266, R268). When these residues were included in our calculation, the *E*_a_ was even lowered to 14.2 kcal/mol. The Arg residues did not significantly affect the distance between C3′ and C8 (3.13 Å vs 3.19 Å, Figure 8A), suggesting that the lower *E*_a_ was achieved by the positive charge or the H-bond interactions with the Arg residues. Among the three Arg residues, R17 had the greatest effect as its removal increased the *E*_a_ by 3.5 kcal/mol (Table S1). The catalytic function of R17 was also tested by activity assays of R17A- and R17K-MoaA. While R17A-MoaA was catalytically inactive, R17K-MoaA exhibited activity comparable to wt-MoaA (Figure SX), suggesting that a positive charge at this position is sufficient for the catalytic function of MoaA. While the auxiliary [4Fe-4S] cluster and the disordered C-terminal tail may also contribute to the transition state stabilization, these results suggested that the conformational constraints and the positive charge provided by the Arg residues are likely important for the catalytic rate acceleration of the 3′,8-cyclization by MoaA.

In all the tested conditions, the free energies of the product were close or slightly higher than those of the reactant. Since the overall transformation of GTP into 3’,8-cH_2_GTP is irreversible, the observation suggests that the addition of GTP C3’• to C8 should be followed by a thermodynamically favorable step. Such a thermodynamic driving force could be provided by the reductive quenching of the aminyl radical. While this step was previously proposed to be catalyzed by an electron transfer from the auxiliary 4Fe-4S cluster^14^, no experimental evidence is currently available. The mechanism of this radical quenching step is currently investigation.

Finally, we also performed DFT calculations using the enol tautomer of GTP C3’• (Table S2). Previously, the coordination of GTP N1 to the auxiliary 4Fe-4S cluster was proposed to stabilize the enol tautomer of GTP^46^. However, in our calculation, in the presence of the three Arg residues, the enol tautomer of GTP C3’• is 22.2 kcal/mol less favored than the keto tautomer (Table S3), while the difference was minimal in the absence of the Arg residues (0.75 kcal/mol). Among the three Arg residues, R266 and R268 had the most significant effects (19.7 kcal/mol, Table S3). Therefore, in the presence of these Arg residues, the keto tautomer is likely favored. In addition, the theoretical activation energy of the 3’,8-cyclization through the enol tautomer was minimally different (1.2 kcal/mol, Table S2) from that with the keto tautomer. While more comprehensive analysis including the auxiliary cluster and the C-terminal tail is needed to identify the predominant tautomer of GTP in the MoaA active site, the above observations suggest that keto-enol tautomerization has minimal to no effects on the 3’,8-cyclization efficiency.

## Discussion

This study describes the comprehensive kinetic characterization of the MoaA-catalyzed reaction. EPR characterization of MoaA reactions revealed that MoaA accumulates an organic radical under turnover conditions. This radical was characterized as 5′-dA-C4′• using a series of isotopically labeled SAM. Intriguingly, these studies revealed that 5′-dA-C4′• carries ^1^H or ^2^H transferred from 3′-position of GTP, suggesting that ^1^H/^2^H-3′ abstraction precedes the 5′-dA-C4′• formation. Subsequent search for shunt products identified the formation of (4′*S*)-5′-dA. Based on these observations, we proposed a presence of a shunt pathway that produces (4′*S*)-5′-dA via 5′-dA-C4′• (Fig. 9). The shunt pathway diverges from the main pathway from GTP-C3′•. While in the main pathway, GTP-C3′• undergoes the intramolecular addition to C-8 to accomplish the 3′,8-cyclization, in the shunt pathway, GTP-C3′• abstracts a H-atom from the 4′-position of 5′-dA, resulting in the formation of 5′-dA-C4′•. The resulting 5′-dA-C4′• is subsequently quenched by abstracting a solvent non-exchangeable hydrogen from a currently unknown hydrogen donor to yield (4′*S*)-5′-dA. Comprehensive kinetic characterization of all the characterized products and 5′-dA-C4′• was also consistent with the proposed model. As described in detail below, the kinetic analysis successfully provided the kinetic constant for the radical-mediated C-C bond formation reaction as 2.7 ± 0.7 s^−1^, which was kinetically unmasked from the preceding rate-determining step (most likely to be the SAM cleavage step) and provided the first evidence that MoaA significantly accelerates this unprecedented 3′,8-cyclization.

**Figure 9.**
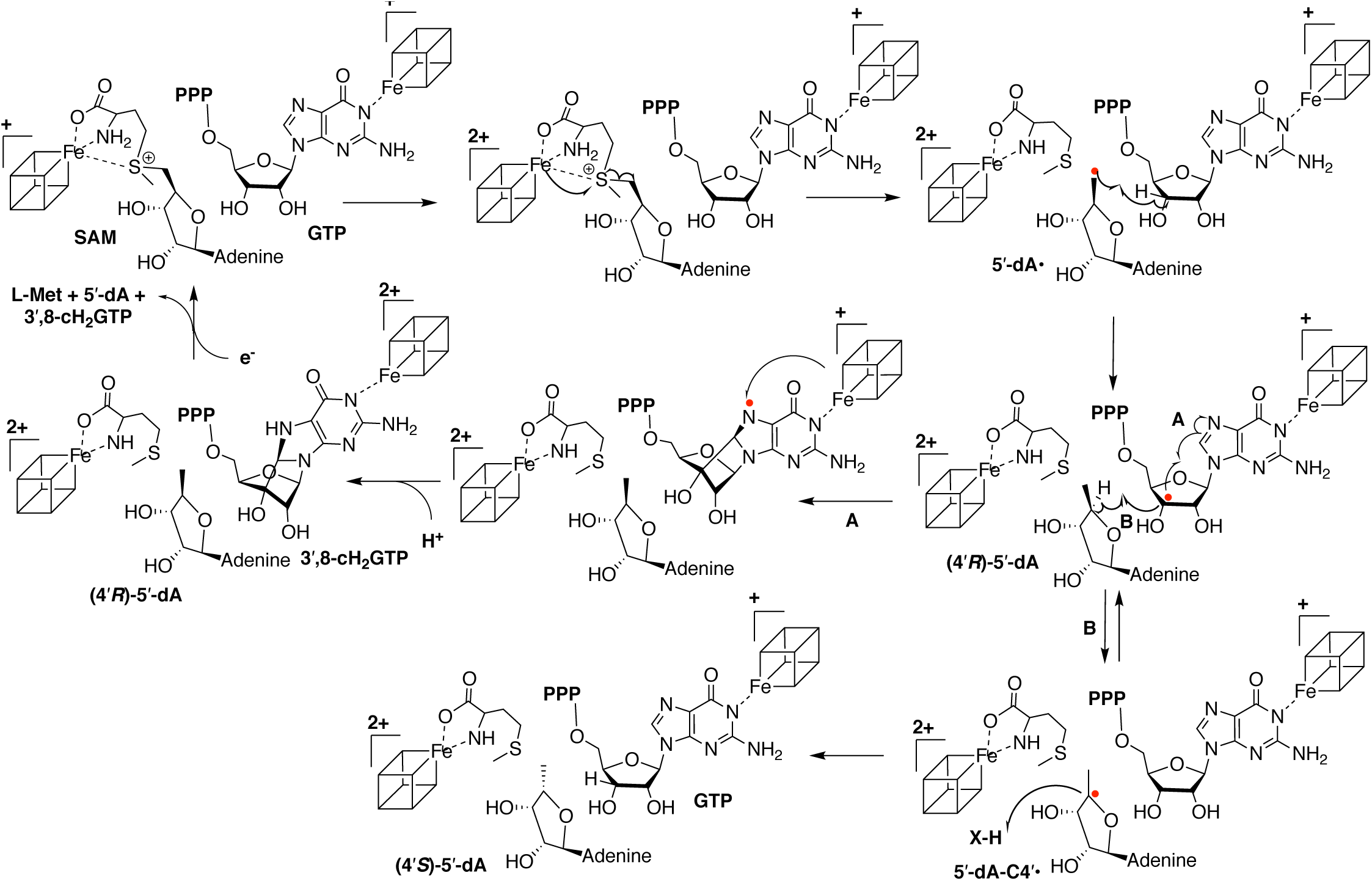
Mechanism of MoaA catalysis with the main pathway that yields 3′,8-cH_2_GTP (path A), and the shunt pathway that produces (4′*S*)-5′-dA via 5′-dA-C4′• (path B). The cubes represent 4Fe-4S clusters.

Kinetically competitive shunt reactions are rare in radical SAM enzymes. Despite the highly reactive nature of radical species, radical SAM enzymes generally catalyze specific chemistry, and rarely catalyzes shunt reactions. Most of the unnatural reactions catalyzed by radical SAM enzymes are reported for reactions with mutant enzymes or unnatural substrates, such as DesII, which is a deaminase on its native substrate TDP-4-amino-4,6-dideoxy-D-glucose while it acts as a dehydrogenase on an unnatural substrate TDP-D-quinovose^47-48^. For wild-type enzymes with natural substrate, the only protein that catalyzes shunt reactions, to our knowledge, is NosL, which catalyzes both the cleavage of Cα-C of L-Trp to produce 3-methyl-2-indolic acid and the cleavage of Cα-Cβ bond of L-Trp to produce 3-methylindole^49-53^. In the current study, MoaA was found as another wild-type radical SAM enzyme with catalytic promiscuity on its natural substrate, where GTP-C3′• can either attack C8 of guanine to form 3′,8-cyclo-GTP aminyl radical or abstracts H-4′ of 5′-dA to form 5′-dA-C4′•.

5′-dA-C4′• is unprecedented in radical SAM enzymes, and likely represents the orientation of 5′-dA relative to GTP-C3′• during the catalysis as well as the kinetics of subsequent 3′,8-cyclization. In a model active site structure of MoaA^14^ (Figure 8B), the distance between GTP C3′ and SAM C4′ is even shorter than the distance between GTP C3′ and SAM C5′ (4.6 Å vs 5.4 Å). Therefore, while this model is likely still missing a significant part of the active site^54^, the analysis is consistent with the abstraction of H-4′ of (4′*R*)-5′-dA by GTP-C3′•. However, this spatial arrangement is not unique among radical SAM enzymes. In the reported crystal structure of BtrN^55^, the distance between C1 of 2-deoxy-*scyllo*-inosamine (DOIA) and SAM C4′ is also comparable to that with SAM C5′ (3.7 Å vs 3.8 Å). However, the accumulation of 5′-dA-C4′• radical has never been observed in other radical SAM enzymes. Thus, the presence of the 5′-dA-C4′• accumulation and the shunt pathway should also reflect the inherently challenging nature of subsequent 3′,8-cyclization. Our kinetic analysis revealed the rate constant *k*_2_ for the 3′,8-cyclization as 2.7 ± 0.7 s^−1^. Although this value is ∼ 2000 times faster than the preceding rate-determining step (k_1_ = 0.0013 ± 0.00002 s^−1^), it is still significantly slower than other radical-mediated intramolecular C-C bond formation reactions in solution (Table 3). The slow 3′,8-cyclization allows GTP-C3′• to abstract H-4′ of 5′-dA that is also slow but comparable to the rate of 3′,8-cyclization (*k*_3_ = 1.9 ± 1.8 s^−1^). Therefore, the structure and the kinetics of MoaA catalysis both contributed to the presence of the shunt pathway, which is likely a compromise of evolving a catalyst for the chemically challenging 3′,8-cyclization.

While the MoaA-catalyzed 3′,8-cyclization is slow compared to other intramolecular radical cyclization reactions (Table 3), it is significantly faster than the theoretical rate of uncatalyzed 3′,8-cyclization. Our analysis suggests that MoaA likely accelerates this reaction by 6 ∼ 9 orders of magnitude, which corresponds to the stabilization of *E*_a_ by 8.5 ∼ 12.4 kcal/mol. This rate acceleration is moderate as an enzyme and comparable to those reported for chorismate mutase^56^. In chorismate mutase, the rate acceleration was proposed to be achieved by orienting the substrate to the near-attack conformation^57-58^ as well as by a transition state stabilization through an electrostatic interaction with an active site Arg residue^59^. Analogously, MoaA may accelerate the 3′,8-cyclization by orienting the GTP conformation optimal for the 3′,8-cyclization. In fact, in our DFT analysis, a decrease of activation energy of ∼ 5.4 kcal/mol was achieved by fixing the structure of GTP-C3′• based on the interactions between GTP and MoaA active site in the crystal and solution structures of MoaA in complex with GTP. In addition, the DFT calculation also suggested that the three catalytically essential Arg residues could also contribute to another ∼ 5 kcal/mol decrease of activation energy through electrostatic or H-bond interaction. Among the three Arg residues, R17 had the most significant effects. Since mutations of these residues in human MoaA homolog have been reported to cause human Moco deficiency disease, understanding of their catalytic functions will not only elucidate the mechanism of MoaA catalysis, but also could provide the foundation for future development of currently incurable human Moco deficiency disease.

Our theoretical analysis also suggested that the addition of GTP C3’• to C8 is slightly endergonic, suggesting that the subsequent aminyl radical reduction needs to be thermodynamically favorable. Although the auxiliary cluster was proposed to serve as an electron donor for this transformation, no experimental evidence is currently available. Even the redox potential or redox state of the auxiliary cluster is not known. Therefore, it is notable that the redox state of the auxiliary cluster could be deduced from the spin relaxation property of 5′-dA-C4′•. Substrate radicals characterized in radical SAM enzymes usually exhibit P_1/2_ ≪ 1 mW at 77 K, a property typical for organic radicals trapped in enzyme active-site without other paramagnetic centers. Thus, the unusually fast relaxation property of the 5′-dA-C4′• signal is unlikely to be caused by the N-terminal radical SAM [4Fe-4S]^2+^ cluster. The [4Fe-4S] clusters from the second subunit of the MoaA dimer is removed by more than 31 Å, and therefore unlikely to cause significant effects on the relaxation properties. Although further investigation is needed, it is most reasonable to conclude that the observed effect is due to the presence of the C-terminal auxiliary cluster in the paramagnetic [4Fe-4S]^1+^ state. If this cluster is in the reduced form during the catalytic cycle, it may then be responsible for the reduction of the putative aminyl radical intermediate. The redox roles have been proposed for auxiliary clusters in the radical SAM enzymes in the SPASM/Twitch family^27, 55, 60^, and the redox potentials have been reported for enzymes under resting states^61^. However, no experimental demonstration has been made about their redox states during the catalytic cycle. Thus, the observations described here represent one of the first experimental evidence for the redox state of auxiliary cluster during the catalytic cycle of any SPASM/Twitch radical SAM enzymes.

In conclusion, we report our comprehensive characterization of MoaA catalysis using EPR and biochemical methods, which revealed a unique shunt reaction that produces (4′*S*)-5′-dA via 5′-dA-C4′•. Detailed kinetic characterization of the shunt and main pathways allowed the determination of the rate constant for the radical mediated C-C bond formation, which is otherwise kinetically masked by the preceding slow SAM cleavage^62^. These results revealed the ability of MoaA to facilitate the chemically challenging 3′,8-cyclization, and provided an important foundation to understand the mechanism of rate acceleration. Future studies of MoaA catalytic mechanisms are important to understand how radical SAM enzymes catalyze chemically challenging radical transformations to provide mechanistic bases on the cause of human Moco deficiency disease.

## Associated content

### Supporting information

Additional experimental details, supplementary methods, and extended data. This material is available free of charge via the Internet at http://pubs.acs.org.

### Notes

The authors declare no competing financial interest.

## Acknowledgement

This work was supported by the Duke University Medical Center and National Institute of General Medical Sciences R01 GM112838 (to K.Y.). We thank Peter Silinski for assistance with the MS measurements. We thank Benjamin G. Bobay at the Duke NMR facility for the assistance in collecting NMR data. EPR spectrometer was supported by an Institutional Development Grant (ID 2014-IDG-1017) from the North Carolina Biotechnology Center.

## Methods

### General

Sodium dithionite (SDT), L-methionine, 2-amino-6,7-dimethyl-4-hydroxypteridine (dimethylpterin, DMPT), 5′-deoxyadenosine (5′-dA), QAE Sephadex A-25 resin were purchased from Sigma-Aldrich, β-mercaptoethanol (βME) from Calbiochem, dithiothreitol (DTT) from Amresco, Guanosine 5′-triphosphate (GTP) from Chem-Impex, Amberlite CG-50 resin from Fisher Scientific, G-25 Sephadex resin from GE Healthcare, Biogel-P2 size exclusion column from Bio-Rad, Ni-NTA agarose resin from Qiagen. [3′-^2^H]GTP was enzymatically synthesized by Dr. Lincoln Scott at Cassia, LLC^31-32^. All the SAM with natural isotope abundance and ^2^H or ^13^C enrichment were enzymatically synthesized from L-methionine and ATP with appropriate isotope labels. *Escherichia coli* (*E. coli*) DH5α and BL21(DE3) competent cells were from Invitrogen. Data plotting and nonlinear least-squares fitting was carried out using GraphPad Prism 7 software. All anaerobic experiments were carried out in an mBraun glovebox maintained at 10 ± 2 °C with an O_2_ concentration < 0.1 ppm. All anaerobic buffers were degassed on a Schlenk line and equilibrated in the glovebox for overnight. All plastic devices were evacuated in the antechamber of glovebox for overnight before use. All HPLC experiments were performed on a Hitachi L-2130 pump equipped with an L-2455 diode array detector, an L-2485 fluorescence detector, an L-2200 autosampler, and an L-2300 column oven maintained at 40 °C. XBridge Amide column (Waters) was used for SAM detection and ODS Hypersil C18 column (Thermo Scientific) for 5′-dA, DMPT, and compound Z. UV-vis absorption spectra were determined using the U-3900 UV-vis ratio recording double-beam spectrometer (Hitachi). EPR spectra were collected using an EMXplus 9.5/2.7 EPR spectrometer system equipped with an In-Cavity Cryo-Free VT system (Bruker Biospin Corporation). Simulation of EPR spectra were performed by EasySpin^66^. Kinetic fitting was performed by Global KinTek Explorer^63-64^. All MS spectra were recorded on a 6224 accurate-mass time-of-flight mass spectrometer equipped with a Dual ESI source, and accurate mass data were obtained by internal calibration using a secondary nebulizer to continuously deliver the reference solution. Flow-injection analysis was conducted in full-scan mode using positive-ion polarity over the range of 80 ∼ 1700 m/z. LCMS analysis was conducted on a PhenKinC18 EVO column (3 × 100 mm) using formic acid in H_2_O/acetonitrile as mobile phase (0 ∼ 100% acetonitrile within 10 min, 0.5 mL/min). NMR samples (isotopically labeled SAM) were prepared in 5 mm diameter NMR tubes with sealed capillary tube containing 0.05 wt. % 3-(trimethylsilyl)propionic-2,2,3,3-*d*_4_ acid, sodium salt in deuterium oxide (99.9 atom % ^2^H). All NMR spectra were recorded on 700 MHz Bruker Advance III NMR spectrometer operated with TopSpin NMR software and analyzed by SpinWorks 4.2.10.

### Expression and purification of recombinant MetK

The plasmids for expression of *E*.*coli* MetK was kindly provided by Dr. Dewey McCafferty. MetK was heterologously expressed in *E. coli* BL21(DE3) cells harboring pET30b-MetK and a pLacI plasmid containing GroEL, GroES chaperon proteins from *S. lividans*^67-69^. Growths were carried out in the presence of kanamycin (35 mg/L) and chloramphenicol (25 mg/L). A single colony was picked and grown in LB medium (5 mL) for 8 hrs followed by dilution into 250 mL LB in a 500 mL baffled flask incubated at 37 °C until the growth reaches saturation (16 hrs). A portion (40 mL) of this culture was used to inoculate 1.5 L of LB medium in a 2.8 L baffled flask and grown at 37 °C until an OD600 reaches 0.6 ∼ 0.8. MetK expression was induced by an addition of 0.25 mM IPTG. The culture continued for an additional 4 hrs. The cells were harvested by centrifugation, frozen in liquid nitrogen, and stored at −80 °C. Typically, 5 ∼ 6 g of wet cell pellet/L of culture were obtained.

Recombinant SAM synthetase, MetK, was aerobically purified. The cell pellets were resuspended and homogenized in 4 volumes of buffer A (50 mM Tris•HCl, pH 7.6, 0.3 M NaCl and 10% glycerol) supplemented with 3 mM βME. The cells were lysed by two passages through a French pressure cell operating at 14,000 psi. The resulting lysate was cleared by centrifugation at 25,000 x *g* and 4 °C for 30 min. The supernatant was loaded onto 20 mL Ni-NTA agarose resin equilibrated in buffer A supplemented with 3 mM βME and 20 mM imidazole. Then the resin was washed with buffer A supplemented with 3 mM βME and 20 mM imidazole for 20 column volumes. MoaC was then eluted using buffer A supplemented with 3 mM βME and 250 mM imidazole. Fractions containing MetK, determined by SDS-PAGE, were exchanged into buffer B (50 mM Tris•HCl (pH 8.0), 100 mM KCl, 10 mM MgCl_2_, and 10% glycerol) using a Sephadex G-25 column. The concentration of MetK was determined based on the UV absorption at 280 nm using the extinction coefficients (ε_280nm_ = 40.1 mM^−1^•cm^−1^) as determined by Edelhoch′s method.

### Enzymatic Synthesis of SAM isotopologues

All natural isotope abundant and the isotopically enriched SAM ([ribose-^13^C_5_]SAM, [5′-^13^C]SAM, [5′-^2^H_2_]SAM, and [4′-^13^C]SAM) were prepared from (isotopically labeled) ATP and L-Met. The isotopically labeled ATP was enzymatically synthesized from isotopically labeled D-ribose or D-glucose using enzymes from de novo purine biosynthesis^31-32^. To enzymatically generate SAM from ATP and L-Met, 30 mL reaction containing 10 mM isotope labeled ATP, 15 mM L-Met, 100 mM Tris•HCl (pH 8.0), 50 mM KCl, 30 mM MgCl_2_, 8% βME and 3 mg/mL MetK was incubated at room temperature and progress was monitored by HPLC. The reaction typically yielded approximately 60% conversion (based on the starting concentration of ATP) after 4 h incubation. The reaction was quenched by adjusting the pH to 5 by adding HCl.

Purification of SAM was based on the previous report^68^. The quenched reaction mixture was cleared by centrifugation, and the supernatant was then loaded onto 100 mL of weak cation exchange resin (Amberlite CG-50) equilibrated with 1 mM sodium acetate pH 5 buffer. The column was washed with the same buffer for 5 column volumes (CV) and then SAM was eluted using 40 mM H_2_SO_4_. Fractions containing SAM determined by UV and HPLC were pooled and titrated with QAE Sephadex A-25 resin in hydroxide form to pH 5. The resultant solution was then concentrated by rotary evaporation to less than 3 mL, and loaded onto a 150 mL Bio-Rad Biogel-P2 size exclusion column equilibrated in H_2_O. SAM was eluted with H_2_O after 2 CV. Fractions containing SAM determined by UV and HPLC were pooled and concentrated by rotary evaporation to less than 3 mL. This solution was degassed and transferred into glovebox for anaerobic assays. The compounds were characterized by MS and NMR and stored at −35 °C in the glovebox. This method generally yields SAM with > 96% purity (analyzed by HPLC) and a total yield of ∼ 25% based on ATP.

### X-band CW-EPR experiments

MoaA assays were carried out under strict anaerobic condition (< 0.1 ppm O_2_). MoaA (450 μM) was first pre-reduced by 1.5 mM sodium dithionite (SDT) in buffer A (100 mM Tris•HCl (pH 7.6), 0.3 M NaCl, 10% glycerol, 5 mM DTT) at 25 °C for 60 min. To initiate the reaction, the pre-reduced MoaA was then mixed with SAM and GTP in buffer A. The final concentration of each component was 300 μM MoaA, 1 mM SDT, 1 mM SAM (or isotopologues) and 1 mM GTP or [3′-^2^H]GTP. For the EPR experiments with [4′-^13^C]SAM or [ribose-^13^C_5_]SAM, 500 μM MoaA was used to improve the signal-to-noise ratio. The samples were transferred into EPR tubes and freeze-quenched manually by submerging in isopentane slush bath (∼ 110 K) in the glovebox. The samples were then taken out of the glovebox and kept in liquid nitrogen for X-band continuous wave EPR characterization.

The EPR spectrum for the [4Fe-4S] clusters were determined at 15 K. EPR parameters were a microwave frequency of 9.36 ∼ 9.39 GHz; a power of 1.589 mW; modulation amplitude of 10 G; modulation frequencies of 100 kHz; time constants of 0.01 ms and a scan time of 100 s. Quantitation of the [4Fe-4S] cluster signal was carried out using a 1 mM Cu(II) sample as standard. The Cu(II) standard was prepared from a ∼ 100 mM CuSO_4_ stock solution in H_2_O with the Cu(II) concentration determined by light absorbance at 810 nm (ε_810nm_ = 11.8 M^−1^•cm^−1^). The stock solution was diluted in 2 M NaClO_4_, 10 mM HCl, 20% glycerol to prepare 1 mM CuSO_4_. EPR spectra of Cu(II) standard were determined under following conditions: A microwave frequency of 9.36∼9.39 GHz; a power of 0.02 mW; modulation amplitude of 4 G; modulation frequencies of 60 kHz; time constants of 0.04 ms and a scan time of 30 s.

The EPR spectra of the organic radical with non-labeled or isotopically labeled substrates were all determined at 45 K, with a microwave frequency of 9.36 ∼ 9.39 GHz; a power of 12.62 mW; modulation amplitude of 2 G; modulation frequencies of 100 kHz; time constants of 0.01 ms and a scan time of 70 s. Quantitation of the organic radical signal was carried out using a 18.0 μM flavosemiquinone radical (Flv•) as standard. Flv• was prepared as previously reported^33-34^. 1.05 eq. of flavodoxin was incubated with 0.5 eq. of SDT at 37 °C for 60 min under anaerobic condition. The concentration of Flv• was determined based on the light absorbance at 579 nm (ε_579nm_ = 4570 M^−1^•cm^−1^). The sample was then transferred into EPR tube and frozen in an isopentane slush bath. EPR parameters for Flv• standard were a microwave frequency of 9.36 ∼ 9.39 GHz; a power of 0.3991 μW; modulation amplitude of 10 G; modulation frequencies of 100 kHz; time constants of 0.01 ms and a scan time of 50 s.

The recorded EPR spectrum was first processed by a baseline correction using Xenon software. For the EPR spectrum of organic radical species, the signal from [4Fe-4S] cluster was subtracted by a control reaction without SAM. The observed signal intensity was determined by calculating the double integration of the first derivative EPR spectrum. This observed intensity was then normalized by the following equation:

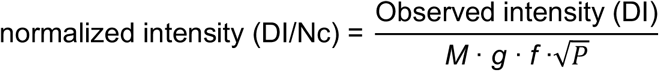

*M* represents modulation amplitude, *P* is the microwave power, *g* is the average *g* value of the radical, *f* is a packing factor related to the inner diameter of EPR tube. The normalized intensity was compared to that of Cu(II) standard to get the exact spin concentration of the sample. All EPR spectral simulations were performed using the EasySpin software^66^.

### Quantitation of (4′*R*)-5′-dA, (4′*S*)-5′-dA and 3′,8-cH_2_GTP by MoaA/MoaC coupled assay

MoaA/MoaC coupled assay was carried out to study the kinetics of the MoaA-catalyzed reaction. MoaA (300 μM) was incubated with MoaC (300 μM), GTP (1 mM), SAM (1 mM), SDT (1 mM) in buffer B (100 mM Tris•HCl (pH 7.6), 0.3 M NaCl, 10% glycerol, 1 mM MgCl_2_, 5 mM DTT). MoaA was first incubated with SDT and MoaC at 25 °C for 30 min to pre-reduce MoaA and transform all the co-purified 3′,8-cH_2_GTP to cPMP. Reaction was then initiated by mixing with GTP and SAM in buffer B. A 30 μL aliquot of the reaction was quenched with 120 μL 3% (w/v) trichloroacetic acid (TCA) at each time point. The quenched sample was centrifuged at 16,100 x *g* for 10 min to remove protein precipitation. To derivatize cPMP into compound Z (CmdZ)^14^, a solution (25 μL) containing 0.2% (w/v) I_2_ and 0.4% (w/v) KI was mixed with 25 μL of the cleared reaction mixture followed by incubation at room temperature for 20 min. The sample was then diluted with mobile phase (0.1% TFA in H_2_O), centrifuged to remove any precipitant, and injected onto the HPLC equipped with an ODS Hypersil column (4.6 × 250 mm). Chromatography was performed by an isocratic elution with 0.1% TFA in H_2_O (0.8 mL/min, 15 min, retention time: 5.5 min) and monitored by fluorescence (excitation: 367 nm, emission: 450 nm). The concentration of CmdZ was quantified based on the standard curve from authentic CmdZ standard. CmdZ standard was prepared by following reported protocol^70^. To quantify the residual unconverted 3′,8-cH_2_GTP, an aliquot of 50 μL quenched sample was chemically derivatized to DMPT by following a previous published method^14^. The quenched mixture (30 μL) was incubated at 95 °C for 5 min to completely hydrolyze MoaA product and release the 6-hydroxy-2,4,5-triaminopyrimidine moiety. After cooling down, the sample was neutralized by 5 μL 0.75 M NaOH and mixed with 15 μL of 0.66% (v/v) 2,3-butanedione in 0.9 M Tris•HCl (pH 8.5). The solution was incubated at 95 °C for 45 min. The resultant solution was then diluted with mobile phase (30 mM ammonium formate, pH 4.5), centrifuged to remove any precipitant, and injected onto the HPLC equipped with an ODS Hypersil column (4.6 × 150 mm). Chromatography was performed by an isocratic elution with 7.5% MeOH, 92.5% 30 mM ammonium formate, pH 4.5 (22.5 min, 1 mL/min, retention time: 11.5 min) and monitored by fluorescence (excitation 365 nm, emission 445 nm). The concentration of DMPT was quantified based on the standard curve of commercial DMPT standard. The amount of 3′,8-cH_2_GTP was calculated as a sum of CmdZ and DMPT. The amount of co-purified 3′,8-cH_2_GTP was subtracted by a MoaA/MoaC reaction with all the components except SAM analyzed by the same protocol. The remaining ∼ 75 μL quenched sample was used for (4′*R*)-5′-dA and (4′*S*)-5′-dA. After removal of precipitation, the quenched sample was directly injected onto the HPLC equipped with an ODS Hypersil column (4.6 × 150 mm). Chromatography was performed by a linear gradient of 0 ∼ 20% MeOH in 30 mM ammonium formate, pH 4.5 (18 min, 1 mL/min, retention time: 12.7 min for (4′*S*)-5′-dA, 14.6 min for (4′*R*)-5′-dA) and monitored by UV absorption at 260 nm. The concentration of both (4′*R*)-5′-dA and (4′*S*)-5′-dA were quantified based on the standard curve of 5′-dA.

### DFT calculation

