## Supplemental PDF for "Accumulation of an unprecedented 5′-deoxyadenos-4′-yl radical unmasks the kinetics of the radical-mediated C-C bond formation step in MoaA catalysis"

**Supplementary Methods**

*Expression, purification and reconstitution of wt-MoaA.*

*Staphylococcus aureus* MoaA with a hexahistidine ($His_6$) tag at the N-terminus was heterologously expressed in *E. coli* BL21(DE3) harboring pET15b-MoaA and a plasmid for the expression of SUF proteins responsible for [4Fe-4S] cluster biosynthesis[1] (kindly provided by Dr. Hänzelmann Schindelin). All cultures were carried out in the presence of chloramphenicol (35 mg/L) and Ampicillin (100 mg/L). A single colony was picked and grown in LB medium (5 mL) for 8 hrs, which was diluted into 250 mL LB in a 500 mL baffled flask incubated at 37 °C with rigorous shaking until the growth reaches saturation (16 hrs). A portion (40 mL) of this culture was used to inoculate 1.5 L of LB medium in a 2.8 L baffled flask, which was incubated at 37 °C with rigorous shaking at 200 rpm until the $OD_{600}$ reaches 0.9~1.0. Then the temperature was lowered to 30 °C and MoaA expression was induced by an addition of 0.5 mM IPTG. The culture was continued for an additional 4 hrs. The cells were harvested by centrifugation, frozen in liquid nitrogen, and stored at -80 °C. Typically, 3~4 g of wet cell pellet/L of culture were obtained.

All purification and reconstitution procedures were carried out anaerobically in m-Braun glovebox (< 0.1 ppm $O_2$, 10 °C). The cell pellet was transferred into the glovebox and resuspended and homogenized in 4 volumes of buffer A (50 mM Tris•HCl, pH 7.6, 0.3 M NaCl and 10% glycerol) supplemented with 3 mM β-mercaptoethanol (βME). The cells were lysed by two passages through a French press cell (> 14000 psi) with the protection of argon. Cellular debris was removed by centrifugation at 25,000 x *g* and 4 °C for 30 min. The supernatant was loaded onto 20 mL Ni-NTA agarose resin (Qiagen) equilibrated in buffer A supplemented with 3 mM βME and 20 mM imidazole. Then the resin was washed with buffer A supplemented with 3 mM βME and 20 mM imidazole for 20 column volumes. MoaA was then eluted using buffer A supplemented with 3 mM βME and 250 mM imidazole. The fractions containing MoaA as judge by Bradford reagent were combined and desalted using a Sephadex G-25 column equilibrated with buffer A supplemented with 5 mM DTT. The concentration of this as-isolated MoaA was quantified by Bradford assay using an apo-wt-MoaA protein as standard. The apo-MoaA was prepared by incubating as-isolated MoaA with 10 mM EDTA and 6 M guanidine•HCl to completely remove [4Fe-4S] cluster. The denatured protein was desalted by G-25 column equilibrated in buffer A supplemented with 5 mM DTT. Any precipitation was removed by centrifugation at 16,100 x *g* and 4 °C for 10 min. Apo-MoaA concentration was determined based on UV absorption at 280 nm using the extinction coefficients ($\varepsilon_{280nm}$ = 26.2 $mM^{-1}\cdot cm^{-1}$) as determined by Edelhoch′s method. The amount of Fe co-purified with MoaA was quantified by Ferrozine assay. Typically, 8 ~ 10 mg as-isolated MoaA/g cell paste was obtained, with 2.5 ± 0.17 eq. of Fe per monomer co-purified.

Reconstitution of the [4Fe-4S] cluster was performed using as-isolated MoaA. To the MoaA solution (diluted to ~ 200 μM), an additional portion of $(NH_4)_2Fe^{II}(SO_4)_2$ and $Na_2S$ was added to reach 9 eq. of $Fe^{2+}$ and $S^{2-}$ per monomer in total (including co-purified Fe). 25 mM stock solution of $(NH_4)_2Fe^{II}(SO_4)_2$ and $Na_2S$ was added dropwisely in 10 cycles over the time course of 30 min. The enzyme was incubated for another 45 min and then buffer exchanged into buffer A supplemented with 5 mM DTT by Sephadex G25 column chromatography. The resulting protein was concentrated to 0.5 ~ 0.7 mM and then flash frozen in liquid nitrogen and stored at -80 °C.

The concentration and amount of Fe were quantified using same protocol as as-isolated MoaA. This reconstitution yielded MoaA with 5.8 ± 0.59 eq. of Fe per monomer (1.45 ± 0.15 eq. reconstituted [4Fe-4S] per monomer). The $k_{cat}$ of the reconstituted wt-MoaA were determined to be 0.097 ± 0.018 $min^{-1}$.

*Expression and purification of wt-MoaC.*

*E. coli* MoaC with a hexahistidine ($His_6$) tag at the N-terminus was expressed in *E. coli* BL21(DE3) harboring pET30b-MoaC. All cultures were carried out in the presence of kanamycin (50 mg/L). A single colony was picked and grown in LB medium (5 mL) for 8 hrs, which was diluted into 250 mL LB in a 500 mL baffled flask incubated at 37 °C with rigorous shaking until the growth reaches saturation (16 hrs). A portion (40 mL) of this culture was used to inoculate 1.5 L of LB medium in a 2.8 L baffled flask, which was incubated at 37 °C with rigorous shaking at 200 rpm until the $OD_{600}$ reaches 0.9~1.0. Then MoaC expression was induced by an addition of 0.5 mM IPTG. The culture was continued for an additional 4 hrs at 37 °C. The cells were harvested by centrifugation, frozen in liquid nitrogen, and stored at -80 °C. Typically, 4 ~ 5 g of wet cell pellet/L of culture were obtained.

*E. coli* MoaC purification was carried out aerobically under 4 °C. The cell pellet was resuspended and homogenized in 4 volumes of buffer A (50 mM Tris•HCl, pH 7.6, 0.3 M NaCl and 10% glycerol) supplemented with 3 mM βME. The cells were lysed by two passages through a French press cell (> 14000 psi) and centrifuged at 25,000 x *g* and 4 °C for 30 min. The supernatant was loaded onto 20 mL Ni-NTA agarose resin (Qiagen) equilibrated in buffer A supplemented with 3 mM βME and 20 mM imidazole. Then the resin was washed with buffer A supplemented with 3 mM βME and 20 mM imidazole for 20 column volumes. MoaC was then eluted using buffer A supplemented with 3 mM βME and 250 mM imidazole. The fractions containing MoaC as judge by Bradford reagent were combined and desalted using a Sephadex G-25 column equilibrated with buffer A supplemented with 5 mM DTT. Protein concentration was determined based on UV absorption at 280 nm using the extinction coefficients ($\varepsilon_{280nm}$ = 7.37 $mM^{-1}$•$cm^{-1}$) as determined by Edelhoch's method. Typically, 10 ~ 12 mg MoaC/g cell paste was obtained. MoaC was then degassed on a Schlenk line using 4 cycles of 30 s evacuation followed by 5 min of argon gas refill and transferred into glovebox for anaerobic assays. The degassed protein was then fast-frozen and stored at -80 °C.

**Table S1. DFT calculation of the free energy of 3',8-cyclization in the keto tautomers.[a]**

| | Reactant (GTP C3'•) | Transition state | Product (3',8-cyclo-GTP aminyl radical) |
|---|---|---|---|
| GTP (no restraints) | 0.00 (3.54) | 24.56 (2.17) | 3.66 (1.55) |
| GTP (with restraints) | 0.00 (3.19) | 19.22 (2.17) | 1.29 (1.55) |
| GTP+R266+R268 (with restraints) | 0.00 (3.15) | 17.72 (2.17) | 1.55 (1.56) |
| GTP+R17+R266+R268 (with restraints) | 0.00 (3.13) | 14.21 (2.16) | -0.36 (1.57) |

[a] Free energy was reported in kcal/mol. Shown in parenthesis are the distances between C3' and C8.

**Table S2. DFT calculation of the free energy of 3’,8-cyclization in the enol tautomers.[a]**

| | **Reactant (GTP C3’•)** | **Transition state** | **Product (3’,8-cyclo-GTP aminyl radical)** |
|---|---|---|---|
| GTP (no restraints) | 0.00 (3.81) | 23.19 (2.18) | 2.42 (1.55) |
| GTP (with restraints) | 0.00 (3.17) | 18.96 (2.18) | 0.63 (1.55) |
| GTP+R266+R268 (with restraints) | 0.00 (3.14) | 16.74 (2.18) | 0.12 (1.56) |
| GTP+R17+R266+R268 (with restraints) | 0.00 (3.12) | 12.97 (2.17) | -2.18 (1.57) |

[a] Free energy was reported in kcal/mol. Shown in parenthesis are the distances between C3’ and C8.

**Table S3. Comparison of the relative free energies of the keto and enol tautomers.**[a]

| | Keto tautomer | | | Enol tautomer | | |
|---|---|---|---|---|---|---|
| | **Reactant** | **TS** | **Product** | **Reactant** | **TS** | **Product** |
| GTP (no restraints) | 0 | 24.56 | 3.66 | 4.25 | 27.44 | 6.67 |
| GTP (with restraints) | 0 | 19.22 | 1.29 | 0.75 | 19.71 | 1.39 |
| GTP+2Arg (with restraints) | 0 | 17.72 | 1.55 | 19.69 | 36.42 | 19.81 |
| GTP+3Arg (with restraints) | 0 | 14.21 | -0.36 | 22.23 | 35.19 | 20.04 |

[a] Free energy was reported in kcal/mol.

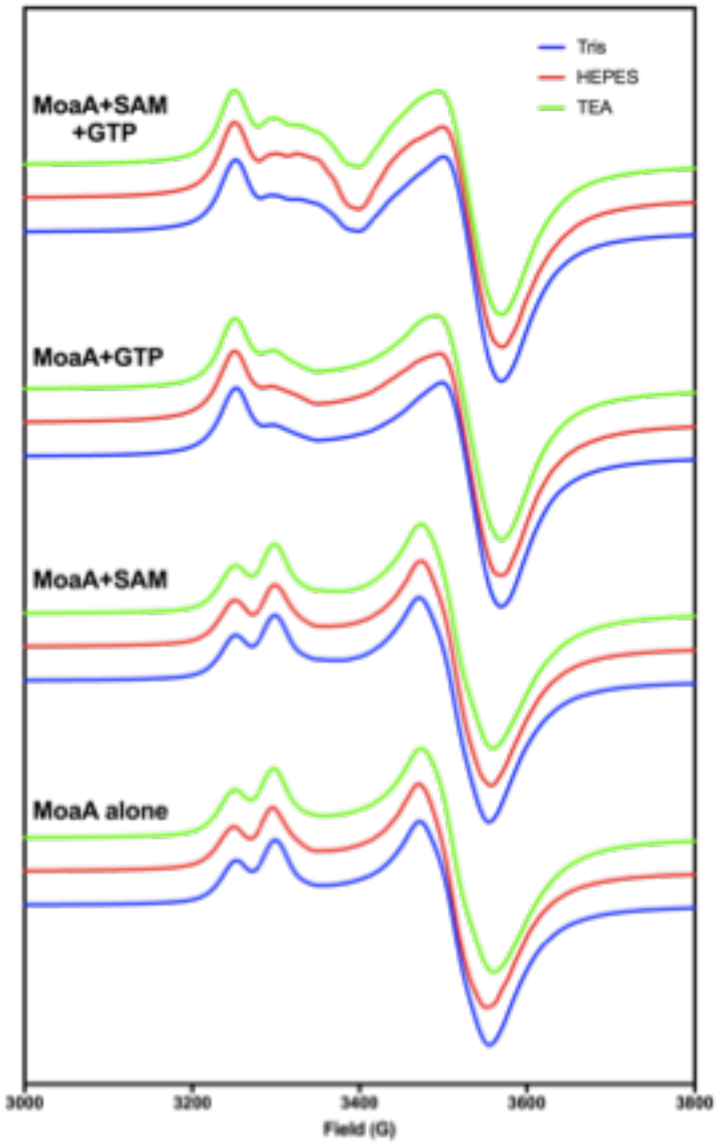

**Figure S1. EPR characterization of MoaA [4Fe-4S] clusters in three different buffers at 15 K.**
Tris (blue traces): 100 mM Tris (pH 7.6), 300 mM NaCl, 10 % glycerol, 5 mM DTT; HEPES (red traces): 50 mM HEPES (pH 7.6), 150 mM NaCl, 10 % glycerol, 5 mM DTT; TEA (green traces): 100 mM triethanolamine (pH 7.6), 300 mM NaCl, 10 % glycerol, 5 mM DTT

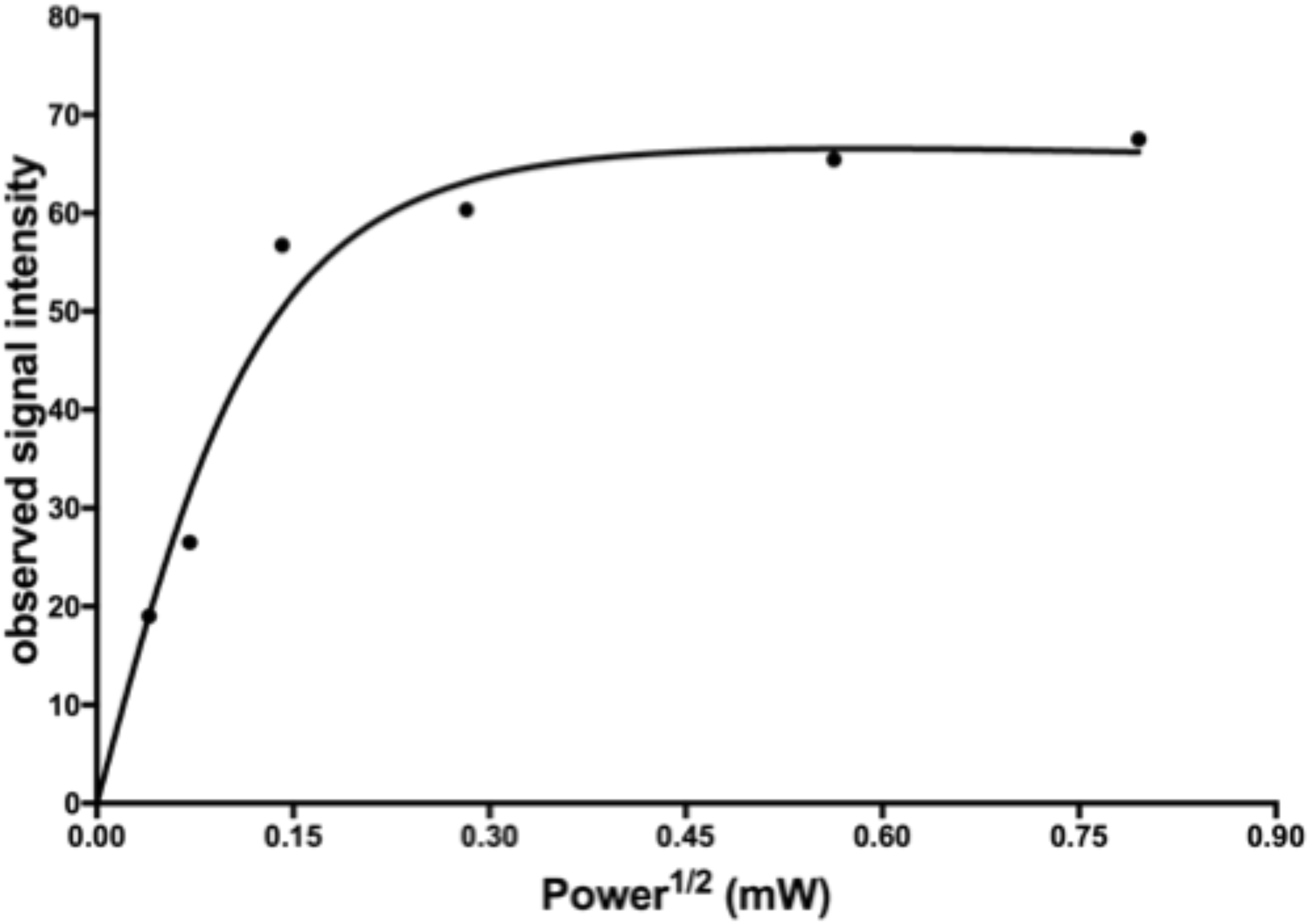

**Figure S2. Relaxation property of the C7′ radical observed in C209A-PolH measured at 77 K[2].**
The solid line represents a non-linear curve fit to eq. 1 by GraphPad Prism with $K$ = 494.9 ± 79.57, $P_{1/2}$ = 23.3 ± 17.7 μW, and $b$ = 1.07 ± 0.18. C209A-PolH was prepared based on previous reported protocol[2].

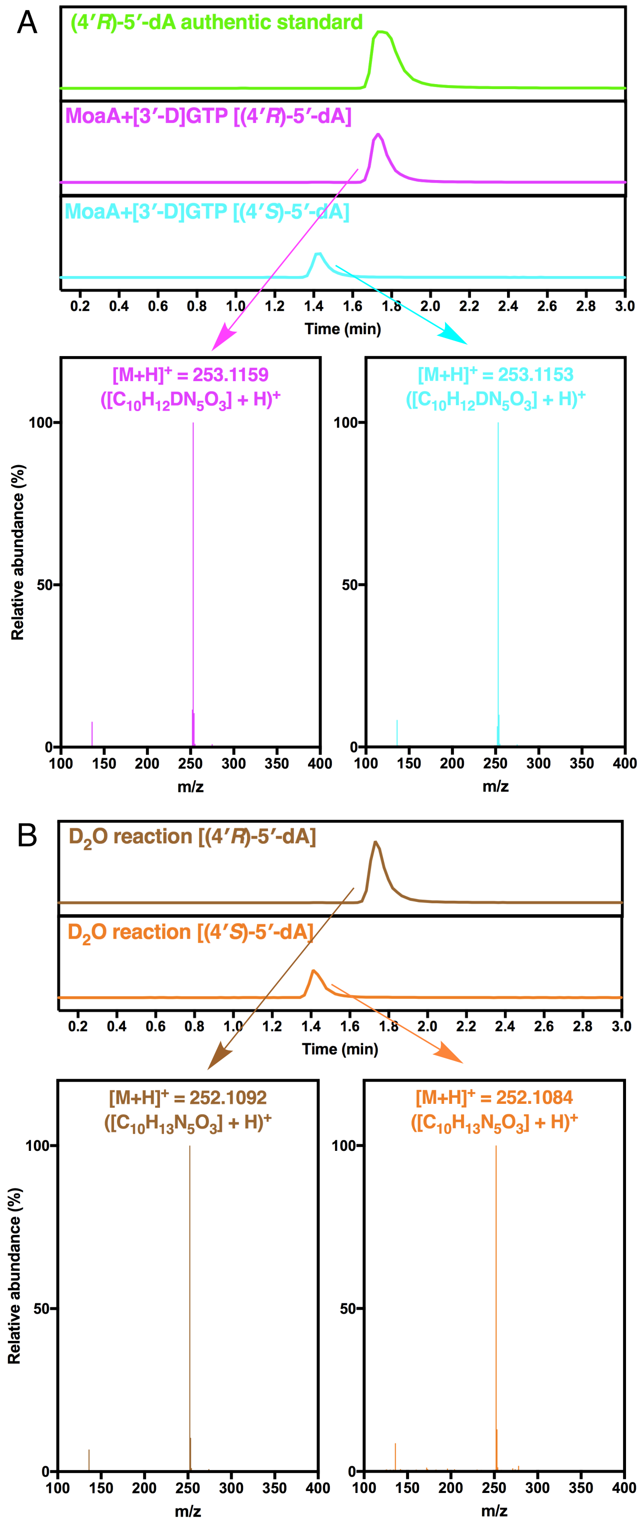

**Figure S3. LCMS characterization of (4′*R*)- and (4′*S*)-5′-dA in MoaA reaction with (A) [3′-D]GTP or (B) in $D_2O$**

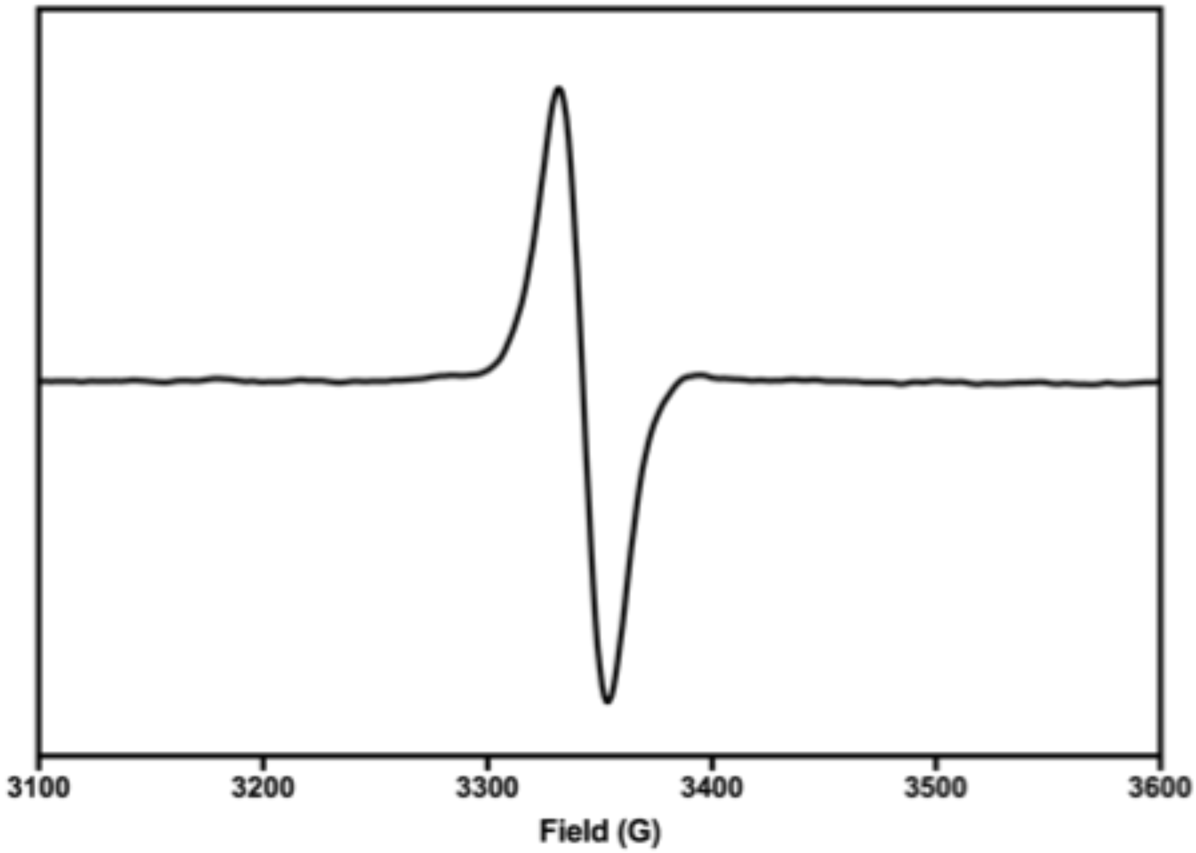

**Figure S4. EPR spectrum of 18.0 μM flavosemiquinone radical at 45 K.**

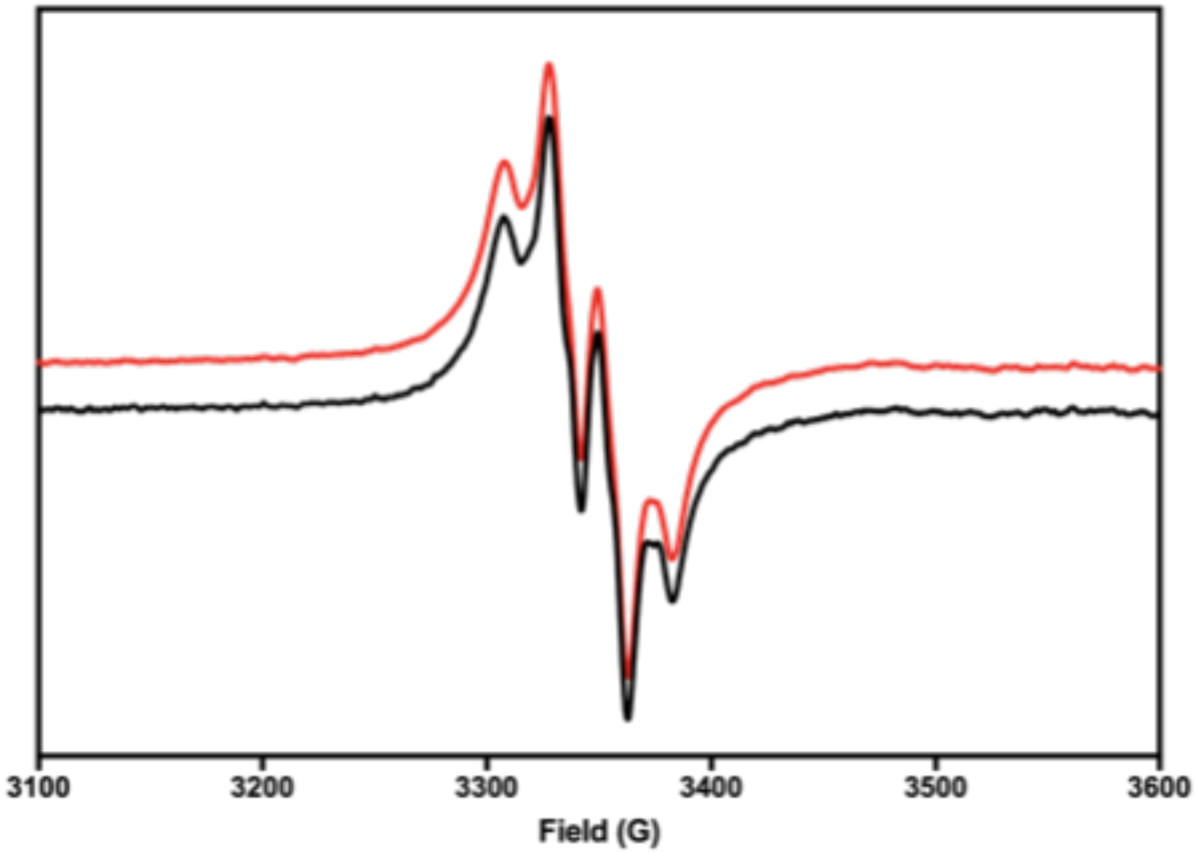

**Figure S5. Comparison of the radical signals in MoaA reaction (black) and MoaA/MoaC coupled reaction (red).**

Both samples were quenched at 2 min in an isopentane slush bath.

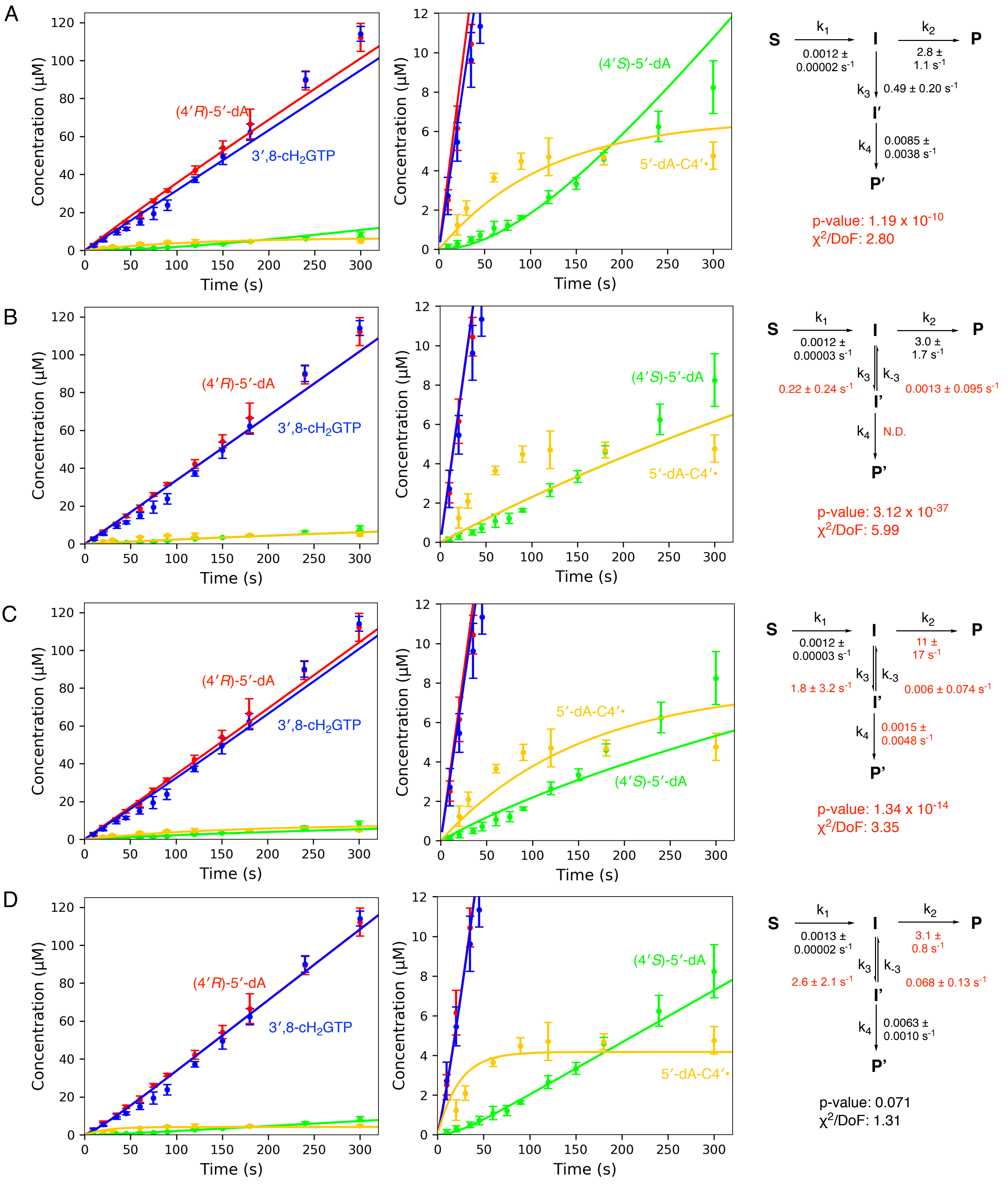
A
Concentration (μM)
Time (s)
(4′R)-5′-dA
3′,8-cH2GTP
(4′S)-5′-dA
5′-dA-C4′•
S
k1
0.0012 ± 0.00002 s-1
I
k2
2.8 ± 1.1 s-1
P
k3
0.49 ± 0.20 s-1
I′
k4
0.0085 ± 0.0038 s-1
P′
p-value: 1.19 x 10-10
χ2/DoF: 2.80
B
Concentration (μM)
Time (s)
(4′R)-5′-dA
3′,8-cH2GTP
(4′S)-5′-dA
5′-dA-C4′•
S
k1
0.0012 ± 0.00003 s-1
I
k2
3.0 ± 1.7 s-1
P
k3
0.22 ± 0.24 s-1
k-3
0.0013 ± 0.095 s-1
I′
k4
N.D.
P′
p-value: 3.12 x 10-37
χ2/DoF: 5.99
C
Concentration (μM)
Time (s)
(4′R)-5′-dA
3′,8-cH2GTP
5′-dA-C4′•
(4′S)-5′-dA
S
k1
0.0012 ± 0.00003 s-1
I
k2
11 ± 17 s-1
P
k3
1.8 ± 3.2 s-1
k-3
0.006 ± 0.074 s-1
I′
k4
0.0015 ± 0.0048 s-1
P′
p-value: 1.34 x 10-14
χ2/DoF: 3.35
D
Concentration (μM)
Time (s)
(4′R)-5′-dA
3′,8-cH2GTP
(4′S)-5′-dA
5′-dA-C4′•
S
k1
0.0013 ± 0.00002 s-1
I
k2
3.1 ± 0.8 s-1
P
k3
2.6 ± 2.1 s-1
k-3
0.068 ± 0.13 s-1
I′
k4
0.0063 ± 0.0010 s-1
P′
p-value: 0.071
χ2/DoF: 1.31

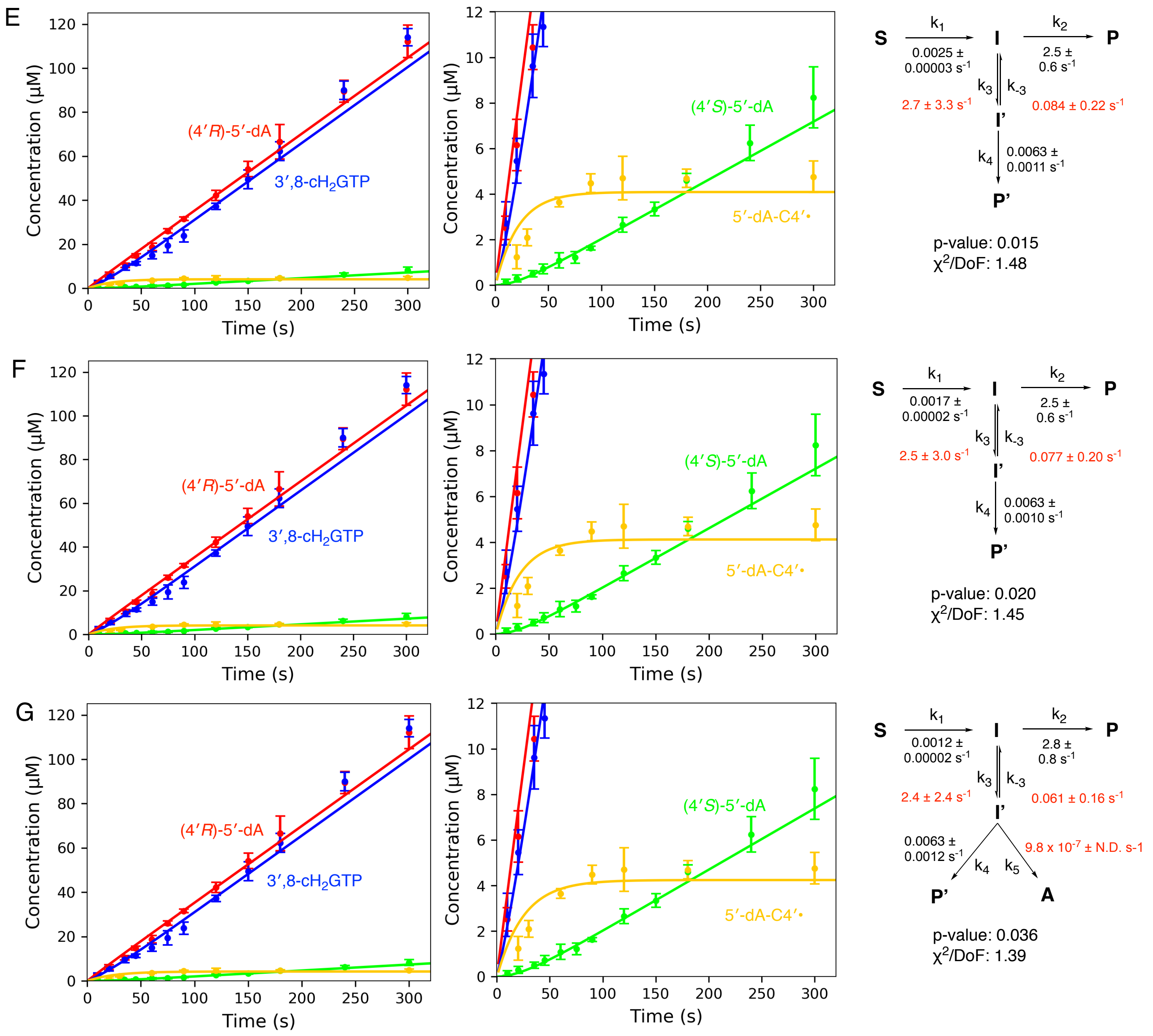

**Figure S6. Kinetic fitting of MoaA catalysis with different models.**

(A) The transformation of **I** to **I’** was set as irreversible. (B, C and D) In these analyses, it was assumed that upon acid quenching, 5′-dA-C4′• was converted to (B) (4′*S*)-5′-dA, (C) both (4′*S*)-5′-dA and (4′*R*)-5′-dA in a 1:1 ratio, or (D) other unidentified molecules. (E and F) The concentration of the Michaelis complex (**S**) were set as (E) 150 μM and (F) 225 μM. (G) In this model, 5′-dA-C4′• was converted to both (4′*S*)-5′-dA and (4′*R*)-5′-dA.

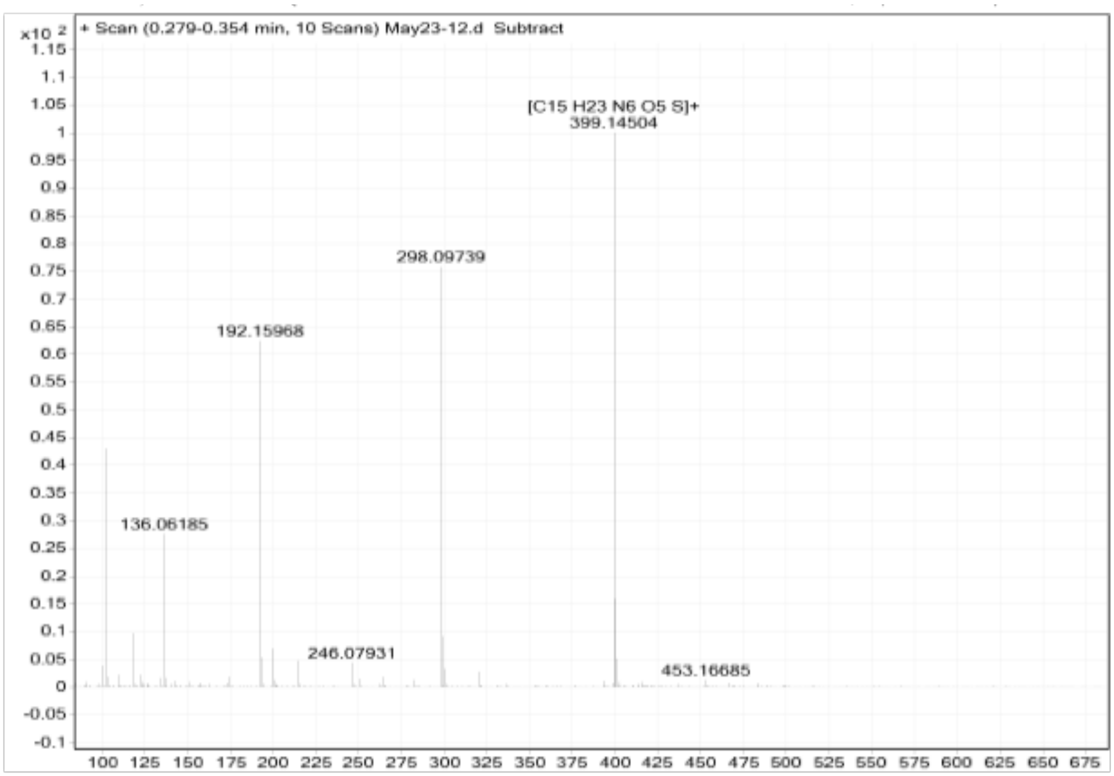

**Figure S7. MS of enzymatically prepared non-labeled SAM.**

*m/z* $[M+H]^+$ calculated for $C_{15}H_{23}N_6O_5S$ 399.1451; found 399.1450.

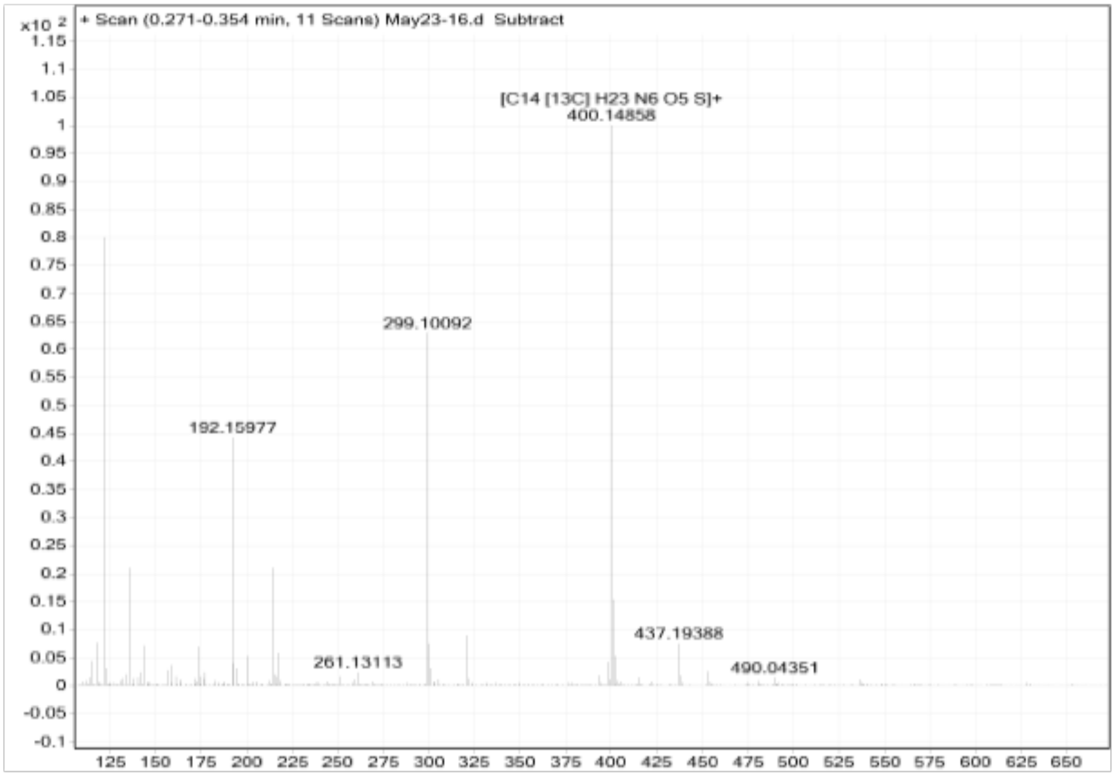

**Figure S8. MS of [5′-$^{13}$C]SAM**

*m/z* $[M+H]^+$ calculated for $C_{14}{}^{13}CH_{23}N_6O_5S$ 400.1484; found 400.1486.

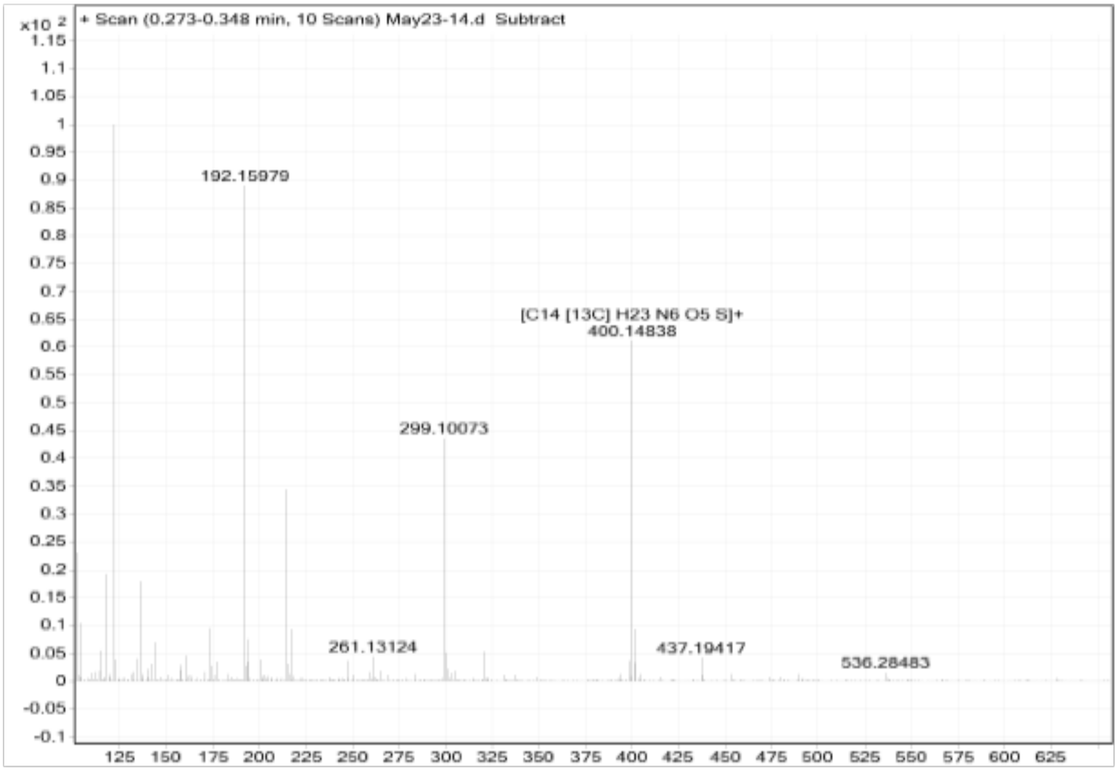

**Figure S9. MS of [4′-$^{13}$C]SAM**

*m/z* $[M+H]^+$ calculated for $C_{14}{}^{13}CH_{23}N_6O_5S$ 400.1484; found 400.1484.

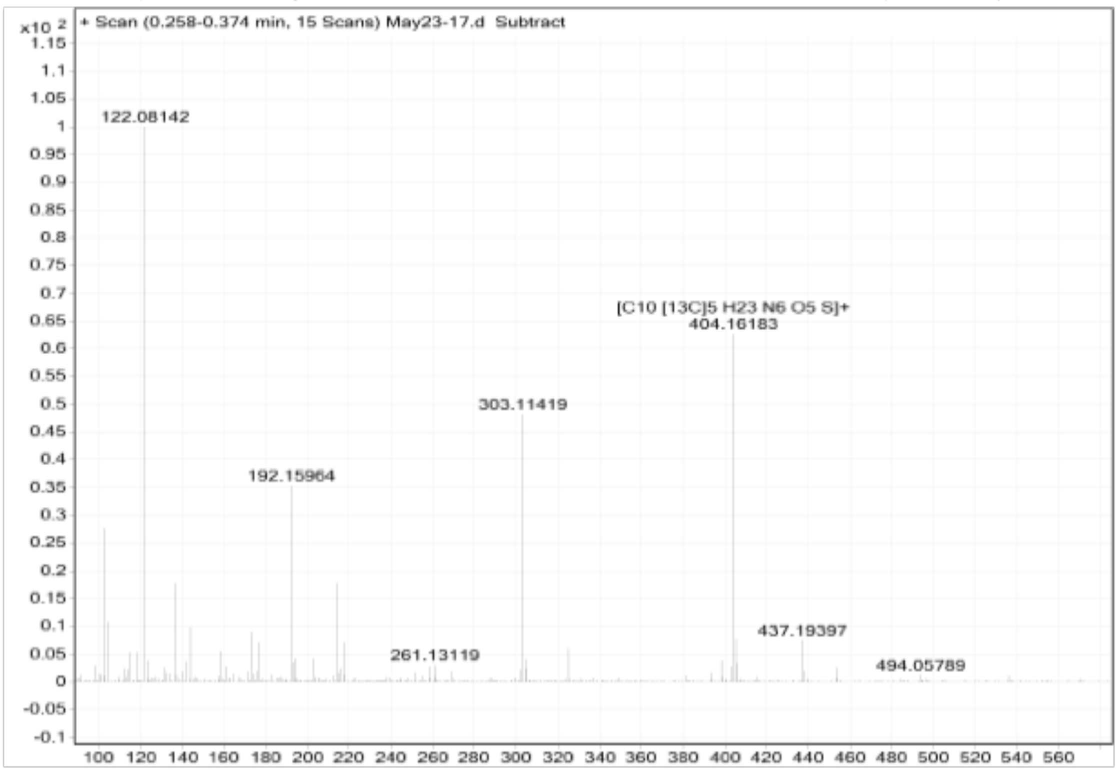

**Figure S10. MS of [ribose-$^{13}C_5$]SAM**

*m/z* $[M+H]^+$ calculated for $C_{10}{}^{13}C_5H_{23}N_6O_5S$ 404.1618; found 404.1618.

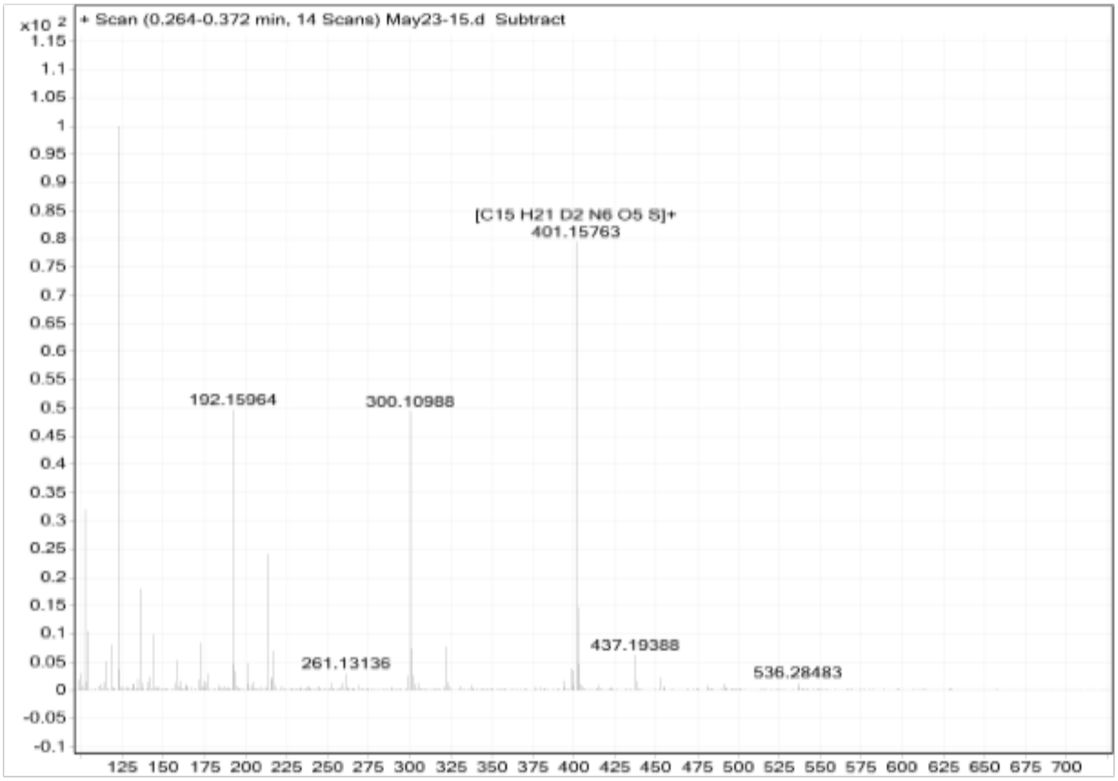

**Figure S11. MS of [5′-$^{2}H_2$]SAM**

*m/z* $[M+H]^+$ calculated for $C_{15}H_{21}{}^{2}H_2N_6O_5S$ 401.1576; found 401.1576.

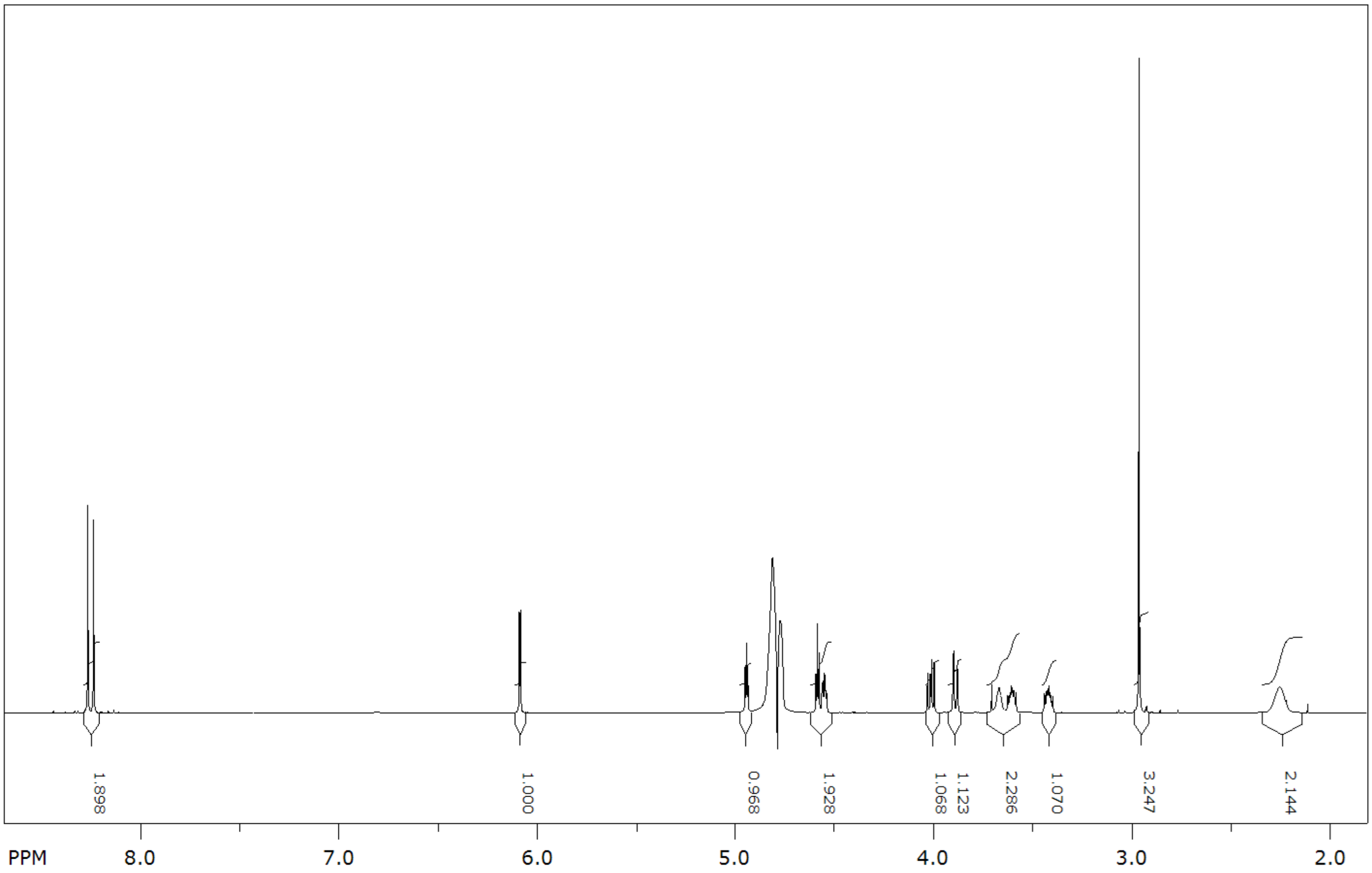

**Figure S12. $^{1}$HNMR of non-labeled SAM**

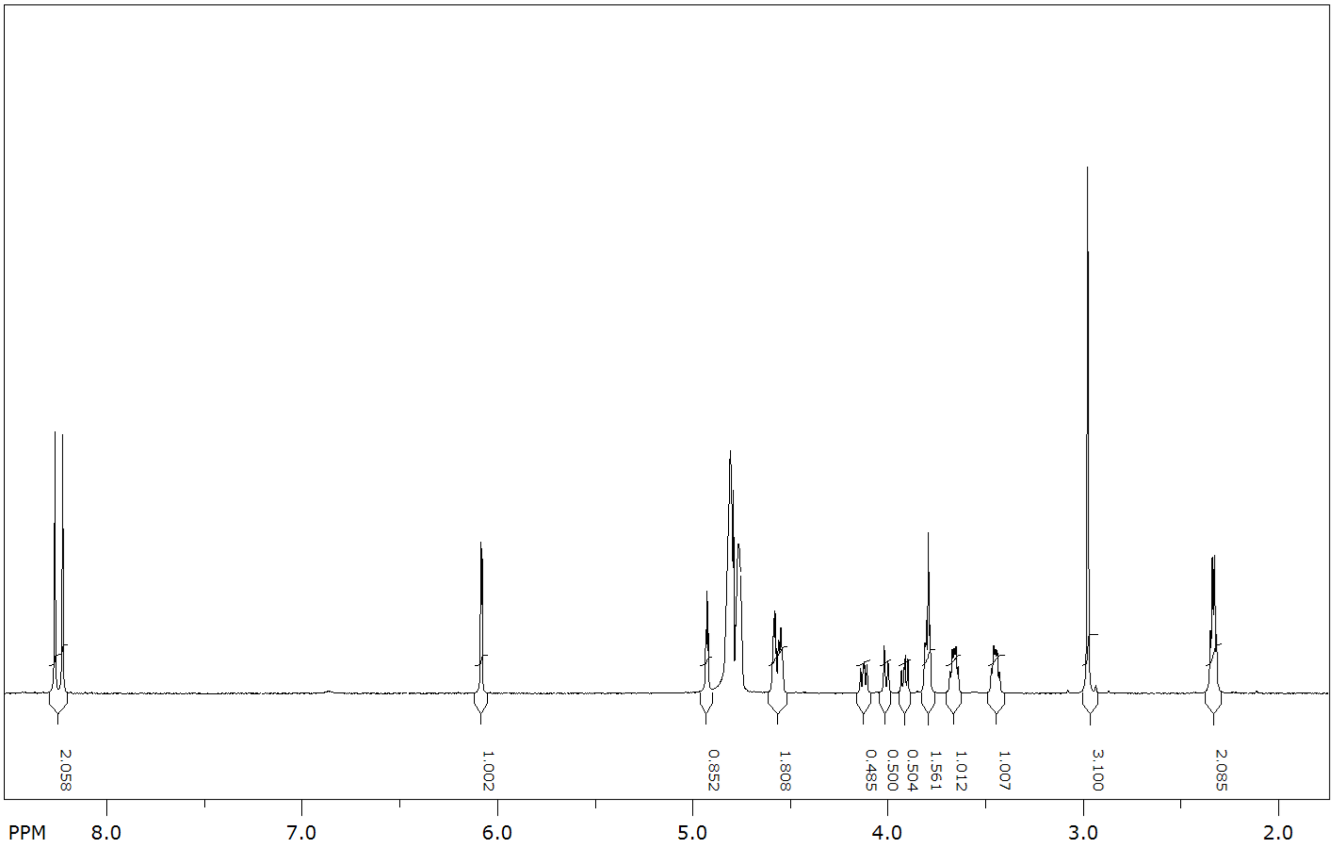

**Figure S13.** **$^{1}$HNMR of [5′-$^{13}$C]SAM**

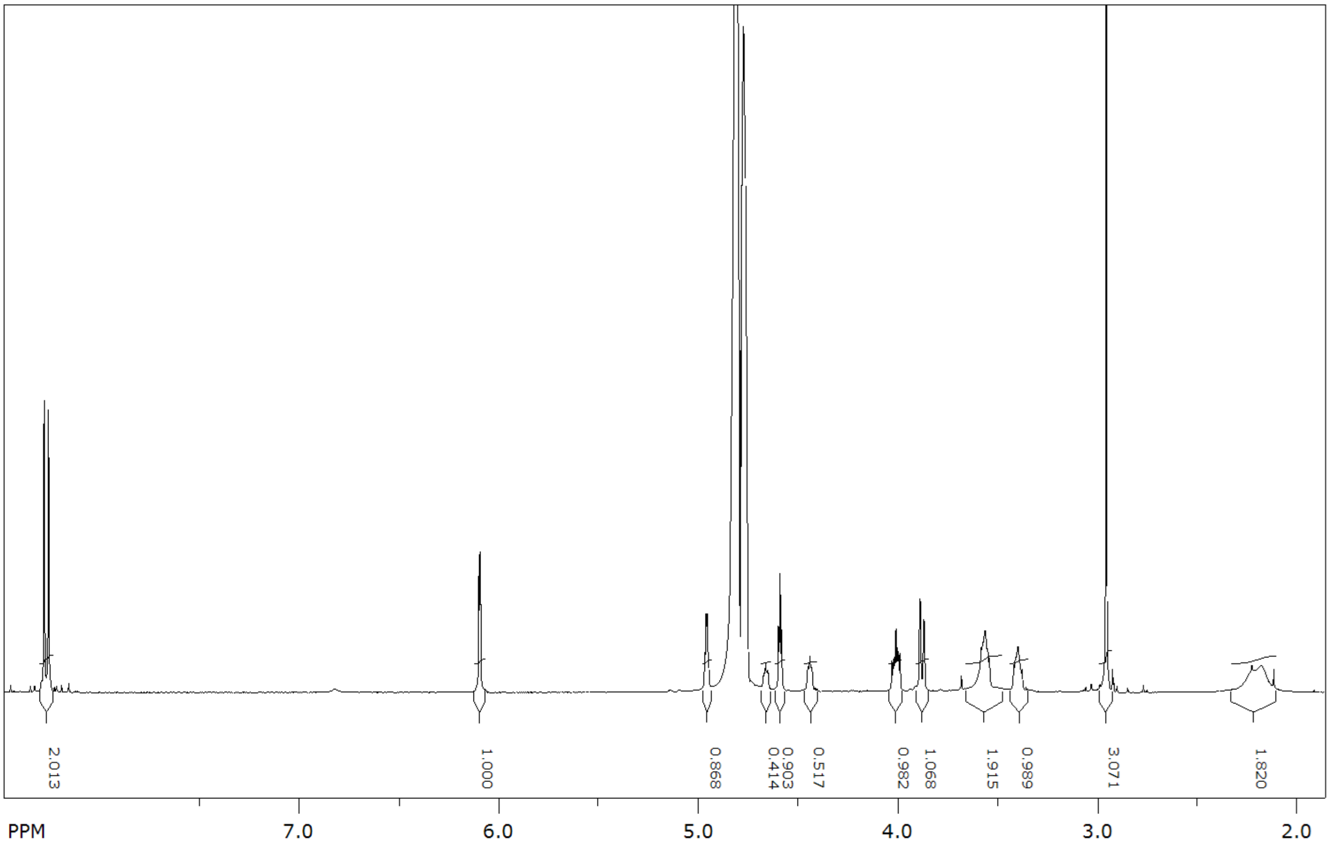

**Figure S14. $^{1}$HNMR of [4′-$^{13}$C]SAM**

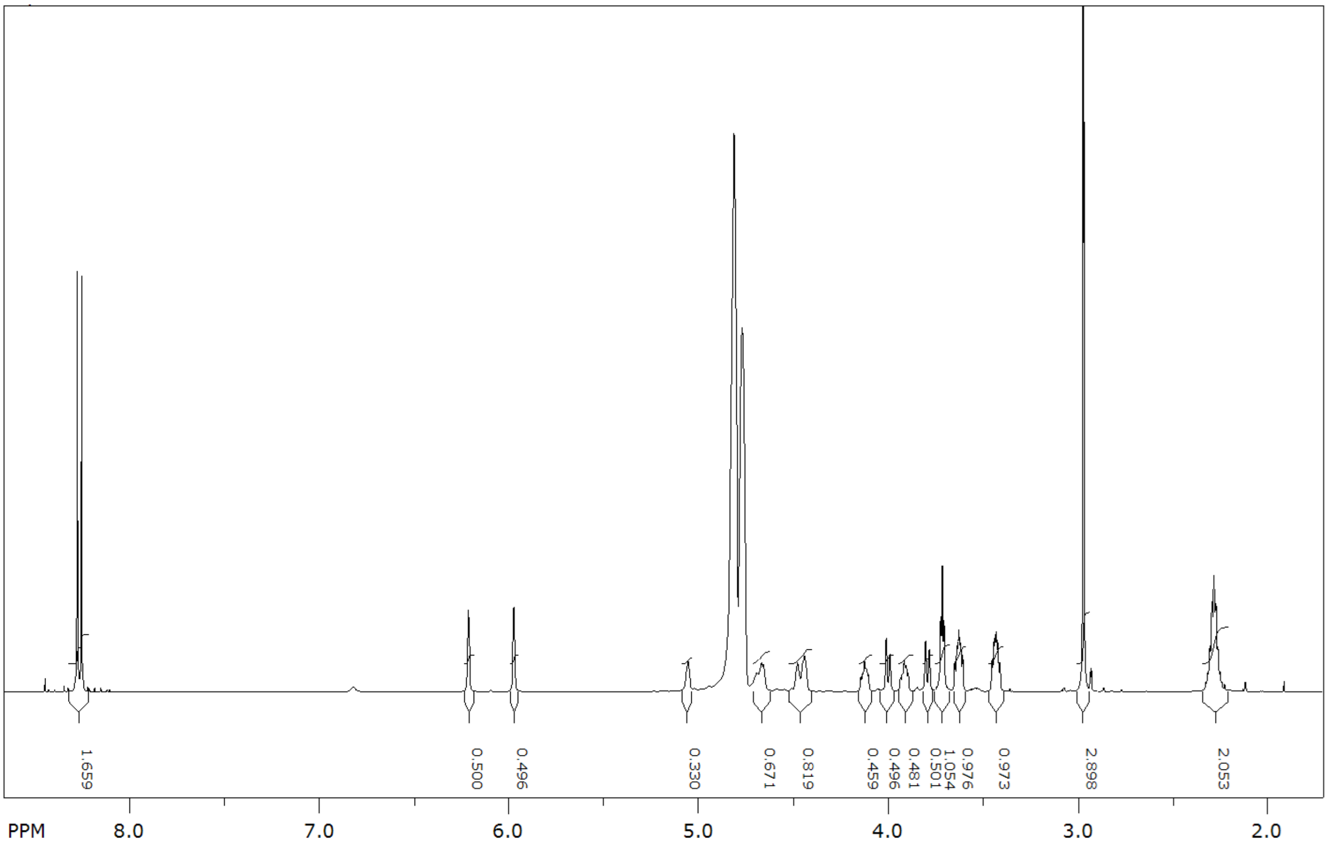

**Figure S15. 1HNMR of [ribose-$^{13}C_5$]SAM**

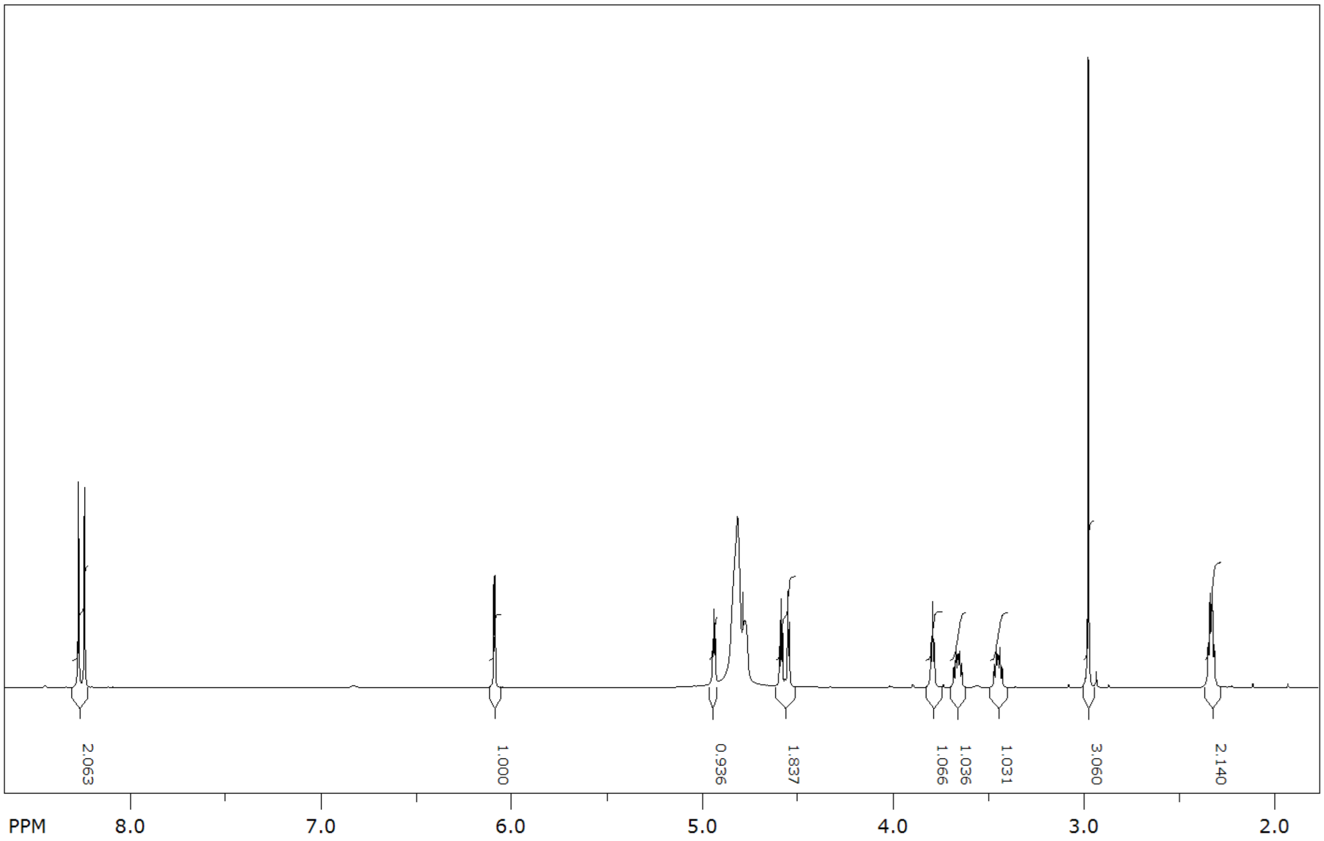

**Figure S16. $^{1}$HNMR of [5′-$^{2}H_2$]SAM**
